# Cancer-specific CTCF binding facilitates oncogenic transcriptional dysregulation

**DOI:** 10.1101/2020.01.17.910687

**Authors:** Celestia Fang, Zhenjia Wang, Cuijuan Han, Stephanie L. Safgren, Kathryn A. Helmin, Emmalee R. Adelman, Kyle P. Eagen, Alexandre Gaspar-Maia, Maria E. Figueroa, Benjamin D. Singer, Aakrosh Ratan, Panagiotis Ntziachristos, Chongzhi Zang

## Abstract

**Background:** The three-dimensional genome organization is critical for gene regulation and can malfunction in diseases like cancer. As a key regulator of genome organization, CCCTC-binding factor (CTCF) has been characterized as a DNA-binding protein with important functions in maintaining the topological structure of chromatin and inducing DNA looping. Among the prolific binding sites in the genome, several events with altered CTCF occupancy have been reported as associated with effects in physiology or disease. However, there is no hitherto a comprehensive survey of genome-wide CTCF binding patterns across different human cancers.

**Results:** To dissect functions of CTCF binding, we systematically analyze over 700 CTCF ChIP-seq profiles across human tissues and cancers and identify cancer-specific CTCF binding patterns in six cancer types. We show that cancer-specific lost and gained CTCF binding events are associated with altered chromatin interactions in patient samples, but not always with DNA methylation changes or sequence mutations. While lost bindings primarily occur near gene promoters, most gained CTCF binding events are induced by oncogenic transcription factors and exhibit enhancer activities. We validate these findings in T-cell acute lymphoblastic leukemia and show that oncogenic NOTCH1 induces specific CTCF binding and they cooperatively activate expression of target genes, indicating transcriptional condensation phenomena.

**Conclusions:** Cancer-specific CTCF binding events are not always associated with DNA methylation changes or mutations, but can be induced by other transcription factors to regulate oncogenic gene expression. Our results substantiate CTCF binding alteration as a functional epigenomic signature of cancer.

## Background

The eukaryotic genome is known to fold into a hierarchical three-dimensional (3D) structure organized by numerous chromatin and transcription factor (TF) proteins[1]. High-throughput technologies such as Hi-C has helped delineate components of 3D genome organization, including topological associating domains (TADs)[2–4] and DNA loops[5]. Studies have shown that various protein factors have roles in chromatin folding that is required for proper gene expression[3,6–9]. One such factor is CCCTC-binding factor (CTCF), a DNA-binding protein that induces chromatin looping and binds at TAD boundaries[10]. CTCF is integral to cell survival as total knockouts in mice are lethal early in embryogenesis and heterozygous knockouts are significantly predisposed to cancer[10–12]. Our previous studies using T-cell acute lymphoblastic leukemia (T-ALL) models have shown that cell-type conserved constitutive CTCF binding sites frequently occur at chromatin domain boundaries and facilitate interactions between TF-bound distal enhancers and their target genes[13]. We demonstrated that TAD boundary intensity associates with CTCF levels, which might also serve to isolate super-enhancers[14]. While CTCF binding at TAD boundaries is usually conserved across diverse cell types and throughout development[15], we and others have shown that CTCF binding within TADs can also exhibit tissue specificity[14,16–18].

The functional importance of CTCF is highlighted in disruptions to CTCF binding coupled with alterations in gene expression, which have been widely observed[19–22]. Deletions of insulator CTCF binding sites can cause aberrant chromatin interactions and differential expression of genes within TADs in developmental disorders and cancers[19,20,23–25]. Many cases of CTCF disruption have been associated with changes in DNA methylation such as in isocitrate dehydrogenase (IDH) mutant gliomas[21], succinate dehydrogenase (SDH)-deficient gastrointestinal stromal tumors (GIST)[22] and immunoglobulin or T-cell receptors[18]. Additionally, the CTCF gene itself or its binding sequence can be mutated and has been suggested to be a haploinsufficient tumor suppressor[12]. CTCF mutations affecting the DNA binding zinc finger domains compromise binding to the genome[26], and can occur in cancer[20,27–29] or abnormal limb development[19]. Mispositioning of even one CTCF binding locus can trigger interactions leading to oncogene activation[20].

While specific CTCF binding sites have been shown to affect gene expression in development, physiology, and cancers[30–34], most others have seemingly little effect on chromatin interaction or gene expression[9, 35]. To date, there is no comprehensive analysis of global and cancer-specific CTCF binding patterns and their functional links to disease-related phenotypes. Here, we performed a novel integrative analysis of a large number of genomic profiles for CTCF as well as other TFs, chromatin marks, gene expression, DNA methylation, somatic mutation, and *in situ* Hi-C to infer CTCF binding site functions. In six different cancer types, we identified cancer-specific gained or lost CTCF binding sites, and showed that gain of CTCF binding in cancer associates with increased chromatin interactions and cancer-specific gene activation, while loss of CTCF binding occurred at promoters of genes present with lower expression in cancer compared to normal cells. We validated our findings in T-ALL, and found that cancer-specific CTCF binding sites are potentially incurred by the activity of oncogenic TFs such as NOTCH1. These findings show that cancers exhibit an oncogenic CTCF binding signature that is intimately tied to chromatin topology and dysregulated gene expression. The putative causative link to oncogenic transcription program suggests that altered CTCF binding is an important component of the mechanism of cancer pathogenesis.

## Results

### Cancers exhibit unique CTCF binding patterns in the genome

CTCF binding sites are among the most stable regulatory elements in the human genome across cell types, compared to gene promoters and distal enhancers (Fig. 1a). To comprehensively study the genomic repertoire of CTCF binding, we collected 771 high-quality CTCF ChIP-seq datasets from the public domain. These datasets cover over 200 human cell types, including normal tissues and multiple cancer types (Fig. S1a, Table S1). We collectively identified 688,429 distinct CTCF binding sites by merging shared peaks called from each dataset (Fig. S1b-c). To study the binding occupancy pattern across cell types, we assigned an occupancy score to each CTCF site by tallying the ChIP-seq datasets exhibiting peaks within the site (Fig. S1a). We obtained a broad spectrum of CTCF binding site distribution from cell-type specific to cross-cell-type conserved (constitutive) (Fig. 1b) and focused on the 285,467 high-confidence CTCF binding sites with occupancy score ≥ 3. We identified 22,097 constitutive CTCF binding sites, which were defined as binding present in at least 80% of all 771 datasets determined by an empirical model (Fig. 1b, S1d).

**Figure 1.**
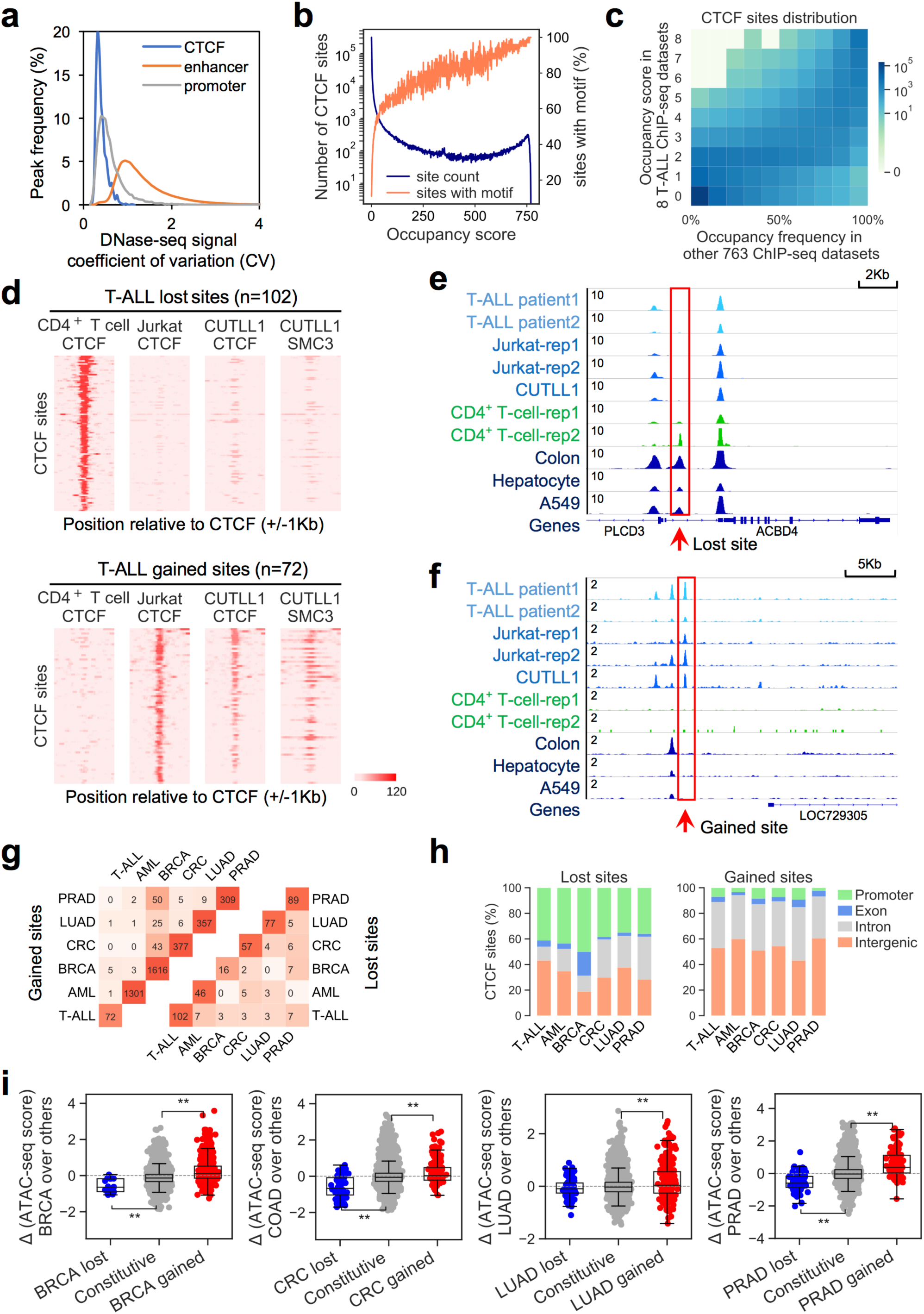
Identification of cancer-specific CTCF binding patterns in the human genome. **a**, Distribution of coefficient of variation of chromatin accessibility at different genomic features, calculated using DNase-seq data from over 60 cell lines from ENCODE. **b**, Distribution of occupancy score for all 688,429 union CTCF binding sites (blue), and percentage of CTCF sites that contain a CTCF motif at each occupancy score (orange). **c**, Distribution of CTCF binding occupancy score in 8 ChIP-seq datasets for T-ALL cell lines (y) and the occupancy frequency score in the other 763 ChIP-seq datasets (x). Color density in each unit represents the number of CTCF binding sites with designated scores. **d**, CTCF ChIP-seq signals at a 2kb region surrounding *T-ALL_lost_* (top) and *T-ALL_gained_* (bottom) binding sites in normal CD4+ T cells and the T-ALL cell lines Jurkat and CUTLL1, and SMC3 signals at the same regions in CUTLL1. **e**, Example of CTCF ChIP-seq signals around a T-ALL-specific lost CTCF binding site. **f**, Example of CTCF ChIP-seq signals around a T-ALL-specific gained CTCF binding site. **g**, Number of identified gained (left) and lost (right) CTCF binding sites in each of the 6 cancer types and number of shared sites between each pair of cancer types. Color density of each element represents the level of similarity measured by Jaccard index. **h**, Genomic distribution of identified lost (left) and gained (right) CTCF binding sites in the 6 cancer types. Promoter regions are defined as +/-2kb from any TSS in the genome. **i**, Differential chromatin accessibility (ATAC-seq) in TCGA patient samples at identified cancer-specific lost (blue), gained (red), and constitutive (grey) CTCF binding sites in each of the 4 cancer types compared to all other TCGA samples. *, p<0.05, **, p<0.001, by unpaired two-tailed Student’s *t*-test.

To identify cancer-specific CTCF binding patterns, we surveyed six cancer types: T-cell acute lymphoblastic leukemia (T-ALL), acute myeloid leukemia (AML), breast cancer (BRCA), colorectal cancer (CRC), lung cancer (LUAD) and prostate cancer (PRAD). These cancers have CTCF ChIP-seq data available in both cell lines and corresponding normal tissues (Table S2). We compared both CTCF occupancy frequencies and normalized CTCF binding levels in samples from each cancer type versus all other datasets (Fig. 1c, S1e-i) as well as their corresponding normal tissue (Fig. S1j) to account for variations due to tissue type specificity. We categorized a CTCF binding event as lost in a cancer if it had a lower occupancy score and lower binding level in the cancer cell lines compared to the corresponding normal tissue and to all other samples. A site was characterized as gained in a cancer if it had a higher occupancy score and higher binding level in the cancer cell lines compared to the corresponding normal tissue as well as to all other samples. Using this approach, we identified lost and gained CTCF binding sites in each of the six cancer types. The CTCF binding patterns at 102 lost and 72 gained sites identified in T-ALL are shown in Fig. 1d-f, comparing normal CD4^+^ T cell with two T-ALL cell lines, Jurkat and CUTLL1[33,34,36–39]. Different cancer types share few commonly lost or gained sites, indicating cancer-type specificity of the identified CTCF binding patterns (Fig. 1g). Across cancer types, however, lost sites are enriched at gene promoter regions (<2kb from any transcription start site, TSS), while gained sites are primarily located at distal and non-coding regions (Fig. 1h). This suggests that different cancers may employ similar mechanisms leading to CTCF binding loss or gain.

We further explored these lost and gained CTCF binding events identified from cancer cell lines in patient samples, to confirm that these unique patterns are not cell line-specific phenomena. In two T-ALL patient samples, 78 of the 102 lost sites (*T-ALL_lost_*) were also depleted in at least one sample, and 33 of the 72 gained sites (*T-ALL_gained_*) are present in at least one sample (Fig. S1k, Table S3). As CTCF binding positively correlates with chromatin accessibility (Fig. S2a, Table S4), we systematically surveyed ATAC-seq data in BRCA, COAD, LUAD and PRAD patient samples from The Cancer Genome Atlas (TCGA)[40], and consistently observed that lost or gained CTCF sites identified from cell lines exhibit lower or higher chromatin accessibility, respectively, compared to the entire TCGA cohort (Fig. 1i), indicating that unique losses or gains in CTCF binding exist extensively in cancer patients. Specific CTCF binding pattern may also be indicative of clinical outcomes, as patients with elevated chromatin accessibility at gained CTCF binding sites have lower overall survival rates (Fig. S2b,c).

### Unique CTCF binding sites link to patterns of chromatin interaction

As CTCF is known to induce DNA looping and is enriched at TAD boundaries[1], we then interrogated the relationship between altered CTCF occupancy and chromatin conformation in cancer. We performed *in situ* Hi-C in Jurkat, healthy donor CD4^+^ T cells, and patient T-ALL cells[5,41,42]. Differential analysis of our Hi-C data reveals that *T-ALL_lost_* sites have decreased and *T-ALL_gained_* sites have increased contact frequencies with their flanking regions (Fig. 2a,b,S3a) compared to constitutive CTCF sites as controls. These findings were corroborated in our two T-ALL patient samples (Fig. 2c-f, S3b,c) and in other malignancies (Fig. 2g,h, S3d, Table S4). Together, these results suggest that cancer-specific alterations to CTCF binding highly associate with changes in local chromatin contacts relative to their normal physiological state.

**Figure 2.**
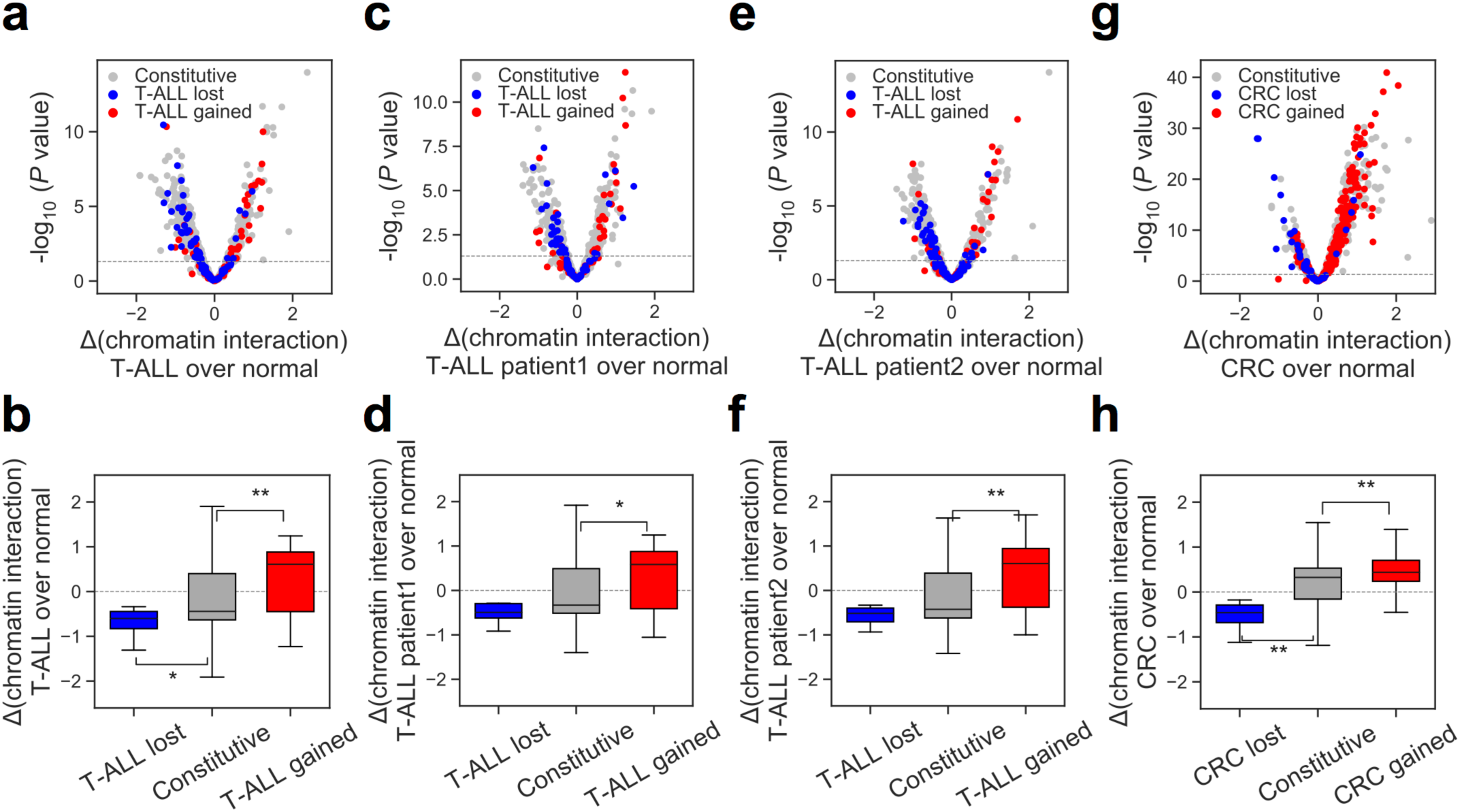
Gained/lost CTCF binding events associate with chromatin dynamics. **a,c,e,g**, Volcano plots showing differential chromatin interaction levels between cancer and normal cells at cancer-specific lost (blue), gained (red), and constitutive (grey) CTCF binding sites, measured by Hi-C. Each point represents the interaction changes between a CTCF binding site and 5kb bins located within 500kb from the site. Horizontal dotted line represents p-value cutoff of 0.05, by paired two-tailed Student’s *t*-test. **b,d,f,g**, Boxplots showing differential interaction frequencies between cancer and normal matched tissues for each group of CTCF binding sites. *, p<0.05, **, p<0.001, by unpaired two-tailed Student’s *t*-test.

In addition to regulating chromatin conformation, CTCF occupancy has been suggested to act as a boundary against spreading of histone modifications between loops and TADs[2, 5]. Therefore, we assessed whether cancer-specific CTCF binding is implicated in histone modification patterns. Using publicly available ChIP-seq data, we examined the activating histone marks H3K4me1, H3K27ac, and the repressive mark H3K27me3, and found that cancer-specific gained CTCF binding associates with increased levels of enhancer marks H3K4me1 and H3K27ac, while lost CTCF sites do not significantly correlate with any of the histone modifications tested (Fig. S4). Analysis of T-ALL patient samples yielded similar results (Fig. S5).

### Cancer-specific CTCF binding gain and loss associate with differential gene expression within chromatin domains

To study the effects of CTCF binding on gene expression, we used an unbiased approach in which we compiled a comprehensive list of all possible combinations of CTCF site and gene pairs that are located within the same chromosome[4,12,13]. We measured both CTCF binding level and gene expression level for each CTCF-gene pair and calculated their correlation across 54 cell types for which both CTCF ChIP-seq and RNA-seq data are available (Table S4). This correlation score represents the association between CTCF binding and gene expression (Fig. 3a). Upon dividing the CTCF-gene pairs into two groups based on whether the paired loci are in the same or different divergently oriented constitutive CTCF-bound chromatin domains[13] (Fig. 3b), we found that pairs in the same domain are more likely to be highly correlated, regardless of genomic distance (Fig. 3c). This indicates that any effect of CTCF binding in regulating gene expression tends to be confined within constitutive CTCF-bound chromatin domains[13]. These domains are highly consistent with TADs identified from Hi-C maps[43] (Fig. S6a,b).

**Figure 3.**
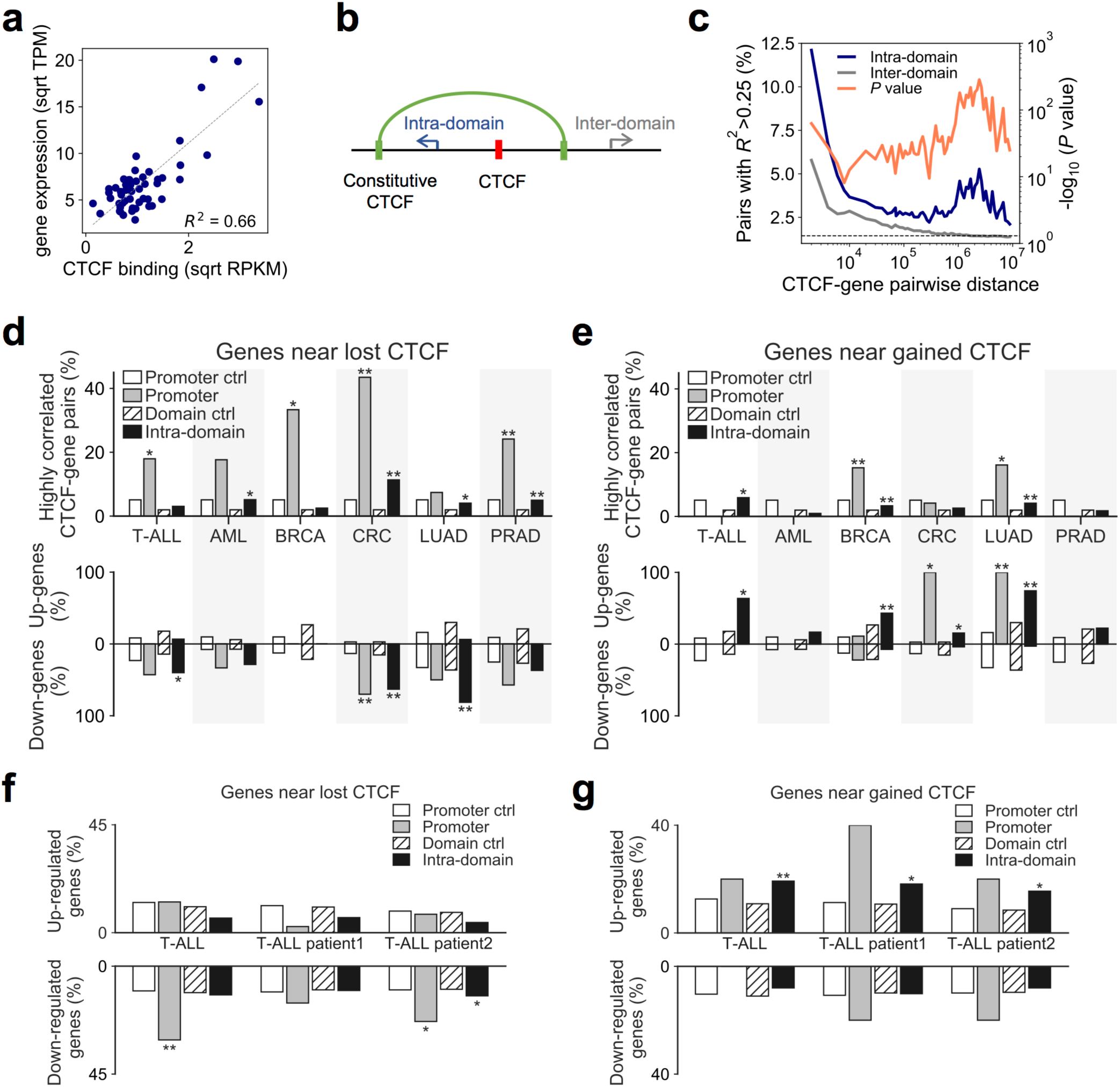
Gained/lost CTCF binding events associate with differential gene expression in cancer. **a**, CTCF ChIP-seq signals (x) and gene expression levels (y) for one CTCF site – gene pair in 54 cell types with matched data available. *R*^2^ is calculated as the association score. Sqrt, Square root; TPM, transcript count per million reads; RPKM, read count per kilobases per million reads. **b**, Schematic of categories of intra-chromatin-domain and inter-chromatin-domain gene-CTCF pairs. **c**, Distribution of highly correlated CTCF-gene pairs (defined as *R*^2^ > 0.25) as a function of the distance between the CTCF binding site and the gene’s TSS. Pairs located within the same CTCF domain (intra-domain, blue) and across different CTCF domains (inter-domain, grey) are plotted separately. *P* values were obtained using two-tailed Fisher’s exact test. **d,e**, Top: Percentage of highly correlated CTCF-gene pairs in which the gene sits within the same domain as a cancer-specific lost (**d**) or gained (**e**) binding site, with constitutive sites as control. “Promoter” designates genes having a promoter region (TSS +/-2kb) that contains its paired CTCF binding site; “Intra-domain” designates genes paired with a CTCF binding site located within the same chromatin domain. Bottom: Percentage of differentially expressed genes (|log2FC|>1, FDR<1e-5) contained within the corresponding group of either Promoter or Intra-domain highly correlated CTCF-gene pairs in the corresponding cancer type. **f,g**, Percentage of genes that are up-regulated (top, log2FC>1, FDR<1e-5) or down-regulated (bottom, log2FC<-1, FDR<1e-5) located in the chromatin domains containing certain group of lost (**f**) or gained (**g**) CTCF sites. *, p<0.05, **, p<0.001, by two-tailed Fisher’s exact test.

We then tested whether those CTCF binding sites specifically lost or gained in cancer associate with expression of genes within the same chromatin domains. If a CTCF binding site is located in a gene promoter region, we directly used that gene as the candidate target. Otherwise, we assigned all genes located within the same domain as the CTCF site as intra-domain candidate target genes. Using this rubric, we found that cancer-specific lost CTCF binding events tend to have higher correlation with their promoter target genes, which are also more likely to be down-regulated in cancer (Fig. 3d). Genes that strongly associate with cancer-specific gained CTCF binding sites, on the other hand, tend to be up-regulated in cancer (Fig. 3e). In general, without considering CTCF-gene pair correlations, genes surrounding lost CTCF binding sites within the same chromatin domain tend to be down regulated while genes surrounding gained CTCF binding sites are more likely to be activated. This relationship holds in both cancer cell lines (Fig. S6c,d) and the two T-ALL patient samples (Fig. 3f,g). Furthermore, we found that genes highly correlated with *BRCA_gained_* intra-domain gained CTCF are enriched for the essential genes identified using CRISPR-screen data from the breast cancer cell line T47D[44], suggesting that gained CTCF is involved in cancer functions (Fig. S6e).

### Differential DNA methylation or sequence mutation do not explain all cancer-specific CTCF binding patterns

We next sought to identify determinants of cancer-specific CTCF binding. To date, the primary identified effectors of variation in CTCF binding at specific loci in cancers include altered DNA methylation at a CTCF motif[20,21,29,45] or mutations affecting the CTCF binding motif[29]. Prior studies have shown that CTCF binding negatively correlates with DNA methylation[15, 46].

We collected DNA methylation profiles from T-ALL, BRCA, CRC, LUAD and PRAD and their corresponding normal tissue, and calculated the differential levels of DNA methylation over a 300bp region centered at each identified lost or gained CTCF binding site. We noticed that some but not all lost CTCF binding sites associate with increased DNA methylation (Fig. 4a), while most gained CTCF sites do not associate with DNA methylation reduction (Fig. 4b). The result was confirmed in the T-ALL patient samples (Fig. S7). We therefore concluded that DNA methylation can only explain a portion of cancer-specific CTCF binding changes.

**Figure 4.**
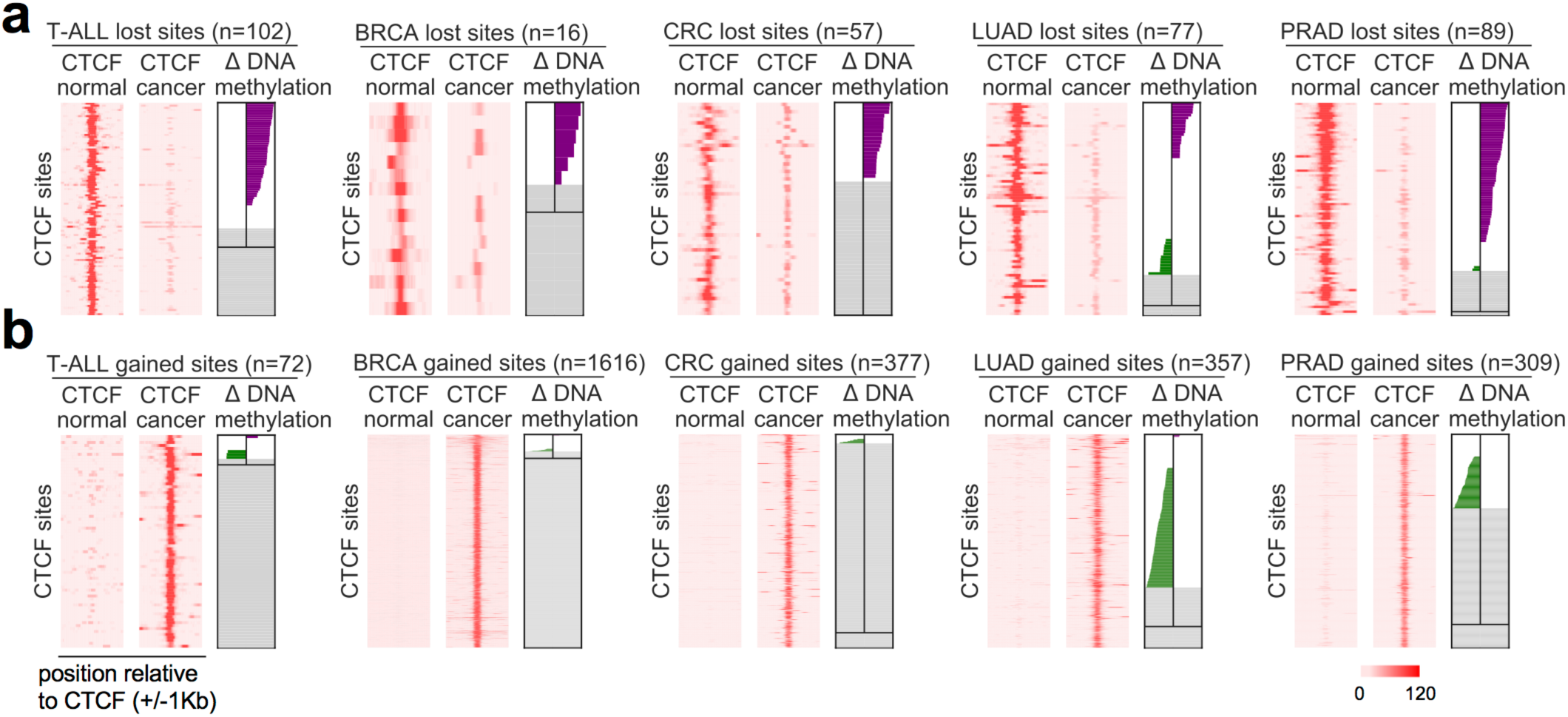
Patterns of differential DNA methylation near cancer-specific lost/gained CTCF sites. **a,b**, ChIP-seq signals and differential DNA methylation levels surrounding specific lost (**a**) or gained (**b**) CTCF binding sites in cancer versus normal matched tissues for each of the 5 cancer types. ChIP-seq heatmaps cover 2kb regions centered at each CTCF site. Differential DNA methylation plots cover 300bp regions centered at each CTCF site. Purple bars represent increased and green bars represent decreased DNA methylation levels (with values in a range from 0 to 100). Grey area with the center vertical line (above the horizontal line) represents regions without enough signal to make confident call of differential methylation. Grey area without the center vertical line (below the horizontal line) represents regions without any detectable methylation signal. Rows in corresponding ChIP-seq and DNA methylation plots are ranked identically.

Stable CTCF binding is highly specific to the presence of its DNA binding motif and can be compromised by mutations affecting the consensus motif sequence[20, 29]. We performed whole genome sequencing (WGS) in T-ALL samples and found very few genetic alterations at gained or lost binding loci (Fig. S8). Using WGS data for AML, BRCA, CRC, LUAD and PRAD patient samples from the International Cancer Genome Consortium (ICGC) [47], we consistently observed that CTCF loss or gain does not associate with mutations altering the consensus binding sequence (Fig. S9). These data show that neither sequence mutations nor DNA methylation changes can fully explain cancer-specific CTCF binding events.

### Cancer-specific gained CTCF co-activates target genes with oncogenic transcription factors

CTCF has been shown to co-bind DNA with other factors to establish DNA loops and control gene expression[19, 48]; thus, we looked for TFs potentially involved in cancer-specific CTCF gain events that associate with dynamic chromatin interaction and increased gene expression. Using our *in situ* Hi-C[4,5,14,49,50] data in T-ALL compared with normal CD4^+^ T cells, we identified genomic regions within the same chromatin domain that interact more frequently with *T-ALL_gained_* CTCF sites (Fig. S10a,b) and used *BART*[51] to identify putative TFs that preferentially bind in these regions. We found that NOTCH1[13], a major oncogenic driver in T-ALL, is one of the top TFs with binding sites enriched in these regions (Fig. 5a). Potential oncogenic TFs in CRC were also identified using the same approach (Fig. S10c, Table S5). Indeed, compared to normal CD4^+^ T cells, gained CTCF sites in T-ALL interact more frequently with “dynamic” NOTCH1 binding sites, previously defined as those sensitive to gamma-secretase inhibitor (γSI) treatment followed by inhibitor washout[13] (Fig. S10d). Furthermore, we analyzed the genome-wide relationship between NOTCH1 and CTCF occupancy in T-ALL and found that both NOTCH1 and dynamic NOTCH1 sites are significantly enriched in chromatin domains containing *T-ALL_gained_* CTCF sites (Fig. 5b, S10e), although NOTCH1 and CTCF do not co-occupy the same loci (Fig. S10f). These *T-ALL_gained_* CTCF sites are also associated with increased levels of H3K27ac in T-ALL, indicative of potential enhancer function (Fig. 5c,d). An example locus is shown in Fig. 5e.

**Figure 5.**
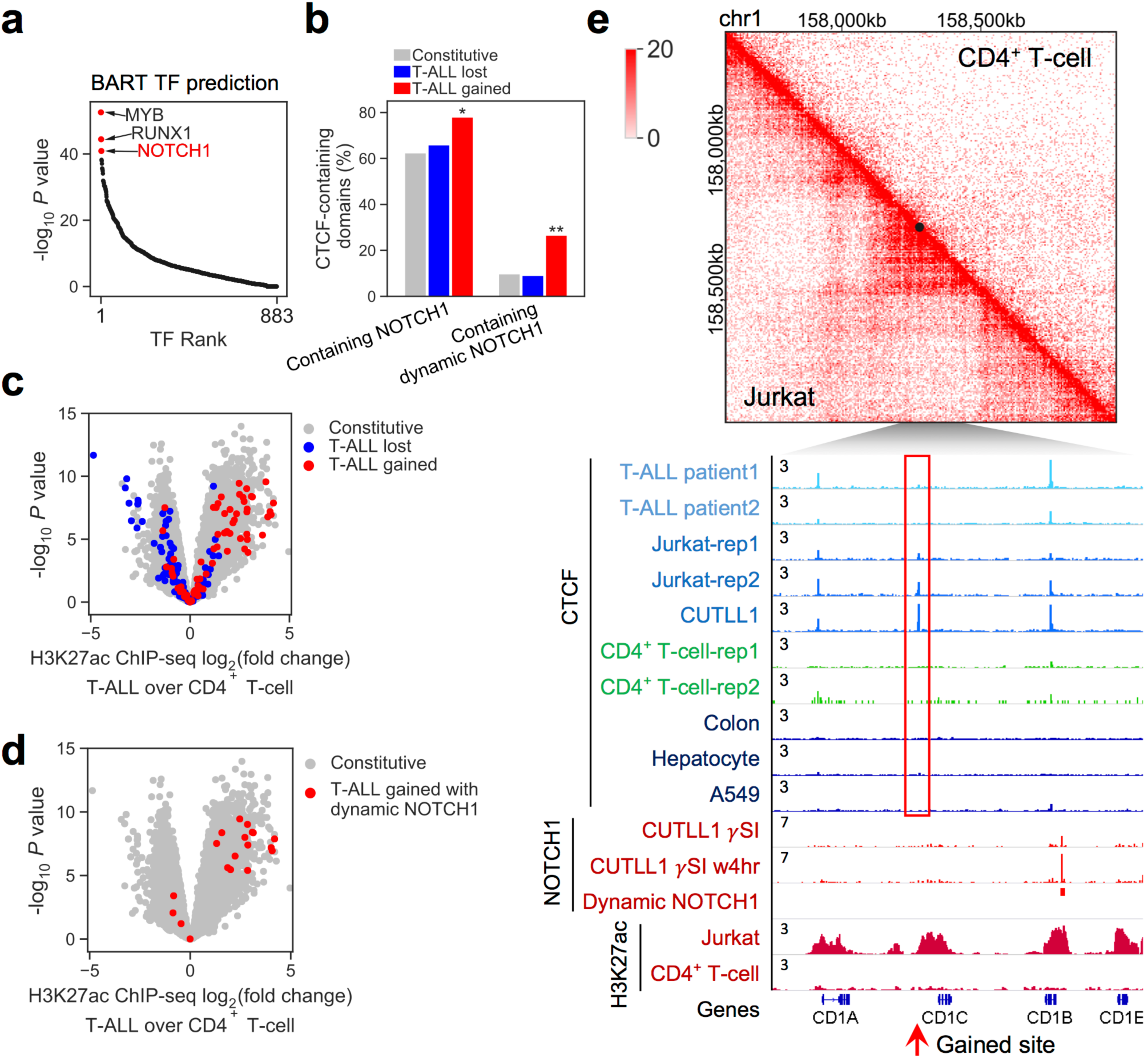
*T-ALL_gained_* CTCF binding associates with oncogenic NOTCH1 binding and increased chromatin interaction. **a**, BART-predicted transcription factors binding in genomic regions that have increased interaction with *T-ALL_gained_* CTCF sites comparing Jurkat cells with normal CD4^+^ T cells. **b**, Percentage of chromatin domains including different groups of CTCF binding that contain a NOTCH1 binding site or a dynamic NOTCH1 binding site. *, p<0.05, **, p<0.001, by two-tailed Fisher’s exact test. **c,d**, *T-ALL_gained_* sites associate with increased H3K27ac level in Jurkat cells. **c**, Volcano plot showing differential H3K27ac level between Jurkat cells and normal CD4^+^ T cells measured by ChIP-seq; each point represents a 10kb region surrounding a CTCF binding site. **d**, Regions containing dynamic NOTCH1 binding sites were highlighted in red. **e**, Example of Hi-C interaction maps and ChIP-seq tracks around a *T-ALL_gained_* CTCF binding site.

### CTCF and NOTCH1 require each other to activate their oncogenic targets in T-ALL

The significant association between *T-ALL_gained_* CTCF binding and dynamic NOTCH1 binding suggests that CTCF might cooperate with NOTCH1 to activate gene expression in T-ALL. To test for dependency of *T-ALL_gained_* CTCF binding on NOTCH1, we treated Jurkat cells with γSI for 72 hours to inhibit the release and nuclear translocation of the intracellular, transcriptionally-active domain of NOTCH1, and then washed out the inhibitor to allow for recovery of intracellular NOTCH1 levels for 16 hours. CTCF ChIP-seq showed that γSI treatment abrogated CTCF binding at most *T-ALL_gained_* sites (Fig. 6a), and a portion of those γSI-sensitive binding events recovered upon γSI washout (Fig. S11a). Meanwhile, chromatin accessibility decreased at *T-ALL_gained_* CTCF sites with γSI treatment compared to DMSO, and significantly reversed after γSI washout (Fig. 6b). These results suggest that functional NOTCH1 binding is required for CTCF binding at *T-ALL_gained_* sites.

**Figure 6.**
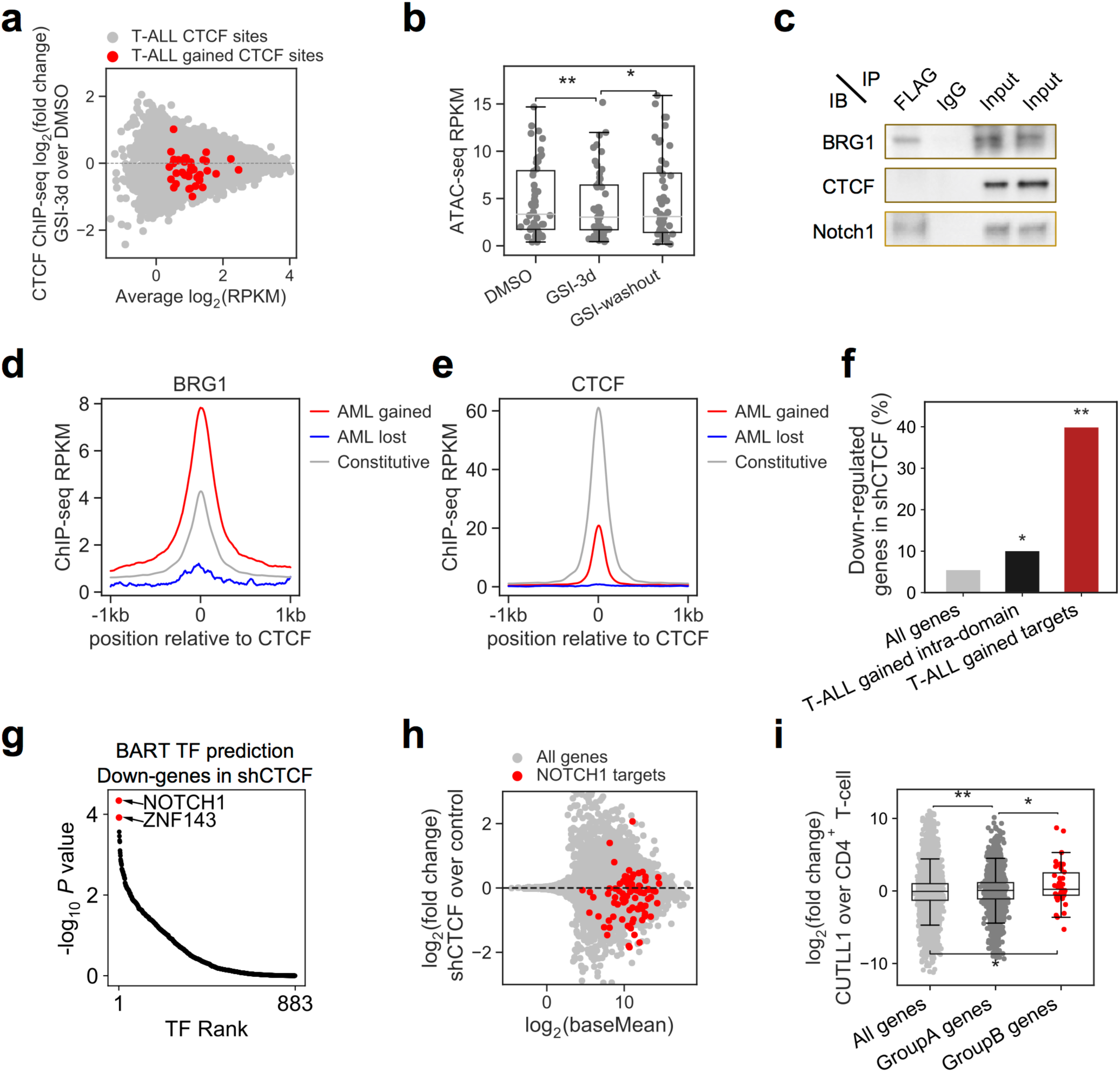
*T-ALL_gained_* CTCF binding facilitates oncogenic NOTCH1 transcriptional activity. **a**, MA plot showing differential CTCF binding levels after GSI treatment in Jurkat cells. Binding levels of most *T-ALL_gained_* CTCF sites (red) are significantly decreased (*P* < 0.001, by unpaired two-tailed Student’s *t*-test). **b**, ATAC-seq levels at *T-ALL_gained_* CTCF sites in Jurkat cells at DMSO, GSI treated for 72 hours, and GSI washout for 16 hours. *, p<0.05, **, p<0.001, by paired two-tailed Student’s *t*-test. **c**, FLAG-NOTCH1 immunopurified proteins from control and NOTCH1-FLAG-expressing CUTLL1 cells were resolved on SDS-PAGE gels and interacting partners are visualized by western blot. IgG was immunopurified as a negative control. IB, immunoblot; IP, immunoprecipitation. **d,e,** ChIP-seq signals for BRG1 (**d**) and CTCF (**e**) surrounding constitutive (grey), *AML_lost_* (blue), and *AML_gained_* (red) CTCF binding sites in AML cell line EOL1. Normalized ChIP-seq read counts (RPKM) covering 2kb regions centered at CTCF binding sites were plotted per 10bp non-overlapped bins. **f,** Percentage of genes in different groups that are down-regulated (log2FC<-0.26, FDR<0.001) in *shCTCF* experiment in CUTLL1. Black: Genes located in the *T-ALL_gained_*-CTCF-containing chromatin domains. Red: Genes located in the *T-ALL_gained_*-CTCF-containing domains that are also up-regulated (log2FC>0.26, FDR<0.001) in T-ALL compared to normal T cell. *, p<0.05, **, p<0.001, by two-tailed Fisher’s exact test. **g**, BART-predicted TFs that target the down-regulated genes (log2FC<-0.58, FDR<0.01) upon *CTCF* silencing experiments in CUTLL1. **h,** MA plot showing differential gene expression after shCTCF treatment in CUTLL1. Most NOTCH1 target genes (red) are downregulated. **i,** Differential gene expression between CUTLL1 and normal T-cells. Group A: genes located in dynamic-NOTCH1-containing domains. Group B: genes located in domains containing both dynamic-NOTCH1 and *T-ALL_gained_* CTCF binding sites. *, p<0.05, **, p<0.001, by unpaired two-tailed Student’s *t*-test.

As NOTCH1 and CTCF do not physically interact with each other (Fig. 6c) and do not co-bind at the same genomic loci (Fig. S10f), we hypothesized that NOTCH1 may mediate the creation of an accessible chromatin configuration to allow for CTCF binding. Recent studies have shown that chromatin remodelers affect CTCF binding[52, 53], and NOTCH1 can interact with the catalytic subunit of the mammalian SWI/SNF chromatin remodeling complex BRG1 (SMARCA4), as well as other members of the BAF and PBAF chromatin remodeling complexes[54]. We confirmed the NOTCH1-BRG1 interaction in our T-ALL cell lines (Fig. 6c), which indicates that NOTCH1 may induce chromatin remodeling. Interestingly, BRG1 in the AML cell lines EOL1 and MOLM13 has higher enrichment at *AML_gained_* CTCF sites than at constitutive CTCF sites (Fig. 6d, S11b)[55], although CTCF itself has lower binding levels at their respective gained sites in both AML and T-ALL than at constitutive sites (Fig. 6e, S11c,d), suggesting that gained CTCF binding might need BRG1 to open chromatin. Thus, a potential mechanism by which NOTCH1 permits *T-ALL_gained_* CTCF binding could occur through BAF complex recruitment to open chromatin for CTCF binding.

The aforementioned findings suggest a potential role for *T-ALL_gained_* CTCF in oncogenic transcription mediated by NOTCH1. To test whether CTCF is required for NOTCH1’s oncogenic transcription function, we knocked down CTCF with short hairpin RNAs (shRNA) in T-ALL cells (CUTLL1). Genes in the same chromatin domains containing *T-ALL_gained_* CTCF binding sites, especially those genes with higher expression in T-ALL compared to normal CD4^+^ T cells were significantly affected by CTCF silencing (Fig. 6f), indicating that *T-ALL_gained_* CTCF sites are the most disrupted in our silencing study. Interestingly, BART analysis revealed that the *shCTCF*-downregulated genes are most likely regulated by NOTCH1 (Fig. 6g). Thus, reducing CTCF levels may disrupt NOTCH1’s ability to activate its target genes. Indeed, most NOTCH1 target genes in CUTLL1 are downregulated in *shCTCF* cells (Fig. 6h). Genes down-regulated in *shCTCF* cells are also significantly enriched for genes down-regulated in γSI-treated cells (Fig. S11e), and reactivated after γSI wash-out (Fig. S11f). These data show that CTCF is required for NOTCH1 to regulate its target genes. Additionally, we found that genes located in chromatin domains containing both dynamic NOTCH1 and *T-ALL_gained_* CTCF sites are most up-regulated in T-ALL compared to normal CD4^+^ T cells (Fig. 6i). Of these T-ALL-upregulated genes, those located in chromatin domains with increased interaction between dynamic NOTCH1 sites and *T-ALL_gained_* CTCF sites are the ones whose expression is the most down-regulated upon CTCF silencing (Fig. S11g). Our collective findings suggest that NOTCH1 and CTCF cooperatively activate oncogenic transcriptional programs in T-ALL.

## Discussion

Through integrative analysis of genomic data collected from the public domain, we presented a comprehensive CTCF binding repertoire in the human genome, from which we identified specific CTCF binding patterns in six distinct cancer types. We characterized a series of genomic and epigenomic features of cancer-specific CTCF binding events using multi-omics profiling techniques including WGS, TF and histone modification ChIP-seq, RNA-seq, ATAC-seq, RRBS, and *in situ* Hi-C. In contrast to previous studies that primarily focused on the effects of mutations or other modifications to CTCF itself or its binding sites[20,21,29,45,56,57], we identified unique CTCF binding patterns in specific cancer types that arise independently of mutations or DNA methylation changes at binding sites. Instead, cancer-specific CTCF recruitment likely results from other TFs that indirectly open chromatin and alter chromatin conformation. CTCF at these sites functions cooperatively with other TFs to facilitate enhancer-promoter interactions and to activate oncogenic transcription programs. In T-ALL, we identified such a cooperative program occurring between NOTCH1 and CTCF, in which NOTCH1 binding is required for gained CTCF binding in the same chromatin domain. This potentially occurs through NOTCH1-induced opening of chromatin at the CTCF binding sites. Gained CTCF binding then cooperates with NOTCH1 to activate transcription of its target genes. Interestingly, we observed substantial enrichment of BRG1 at gained CTCF binding sites (Fig. 6d), as well as a direct protein-protein interaction between NOTCH1 and BRG1 (Fig. 6c). Although previous studies suggested that CTCF and BRG1 might physically interact^57^, we do not find this to be the case in T-ALL (Fig. S11h).

The dynamic interactions involving multiple factors and novel CTCF binding within a single chromatin domain may indicate the formation of phase-separated transcriptional condensates at super-enhancers[58–60]. In T-ALL, NOTCH1 binding drives the establishment of super-enhancers[13]. Thus, *T-ALL_gained_* CTCF binding may be recruited by clusters of TFs and co-activators including chromatin remodeling complexes within phase-separated transcriptional condensates around super-enhancers. The potential for NOTCH1 as a master TF to direct the formation of 3D spatial clusters has been reported recently[61]. Transcriptional condensates maintain a highly active environment, which is consistent with the enrichment of H3K27ac observed near *T-ALL_gained_* CTCF sites (Fig. 5c). By inducing the frequency of chromatin contacts, gained CTCF binding may function to maintain the condensation state that helps drive transcription. A schematic model of the relationships between dynamic NOTCH1 binding, CTCF gain and activation of NOTCH target genes in T-ALL is shown in Fig. 7.

**Figure 7.**
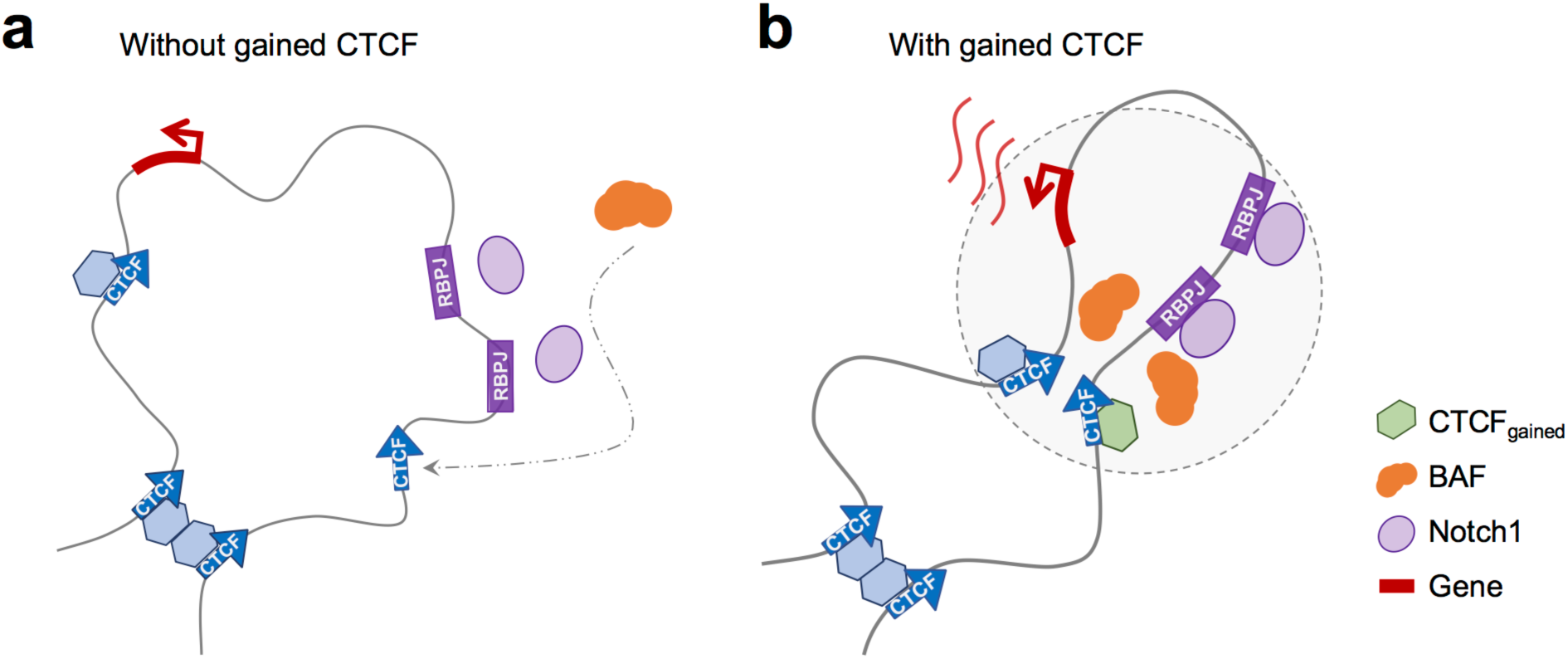
Schematic model of CTCF facilitated oncogenic transcriptional activation in T-ALL. **a**, Without gained CTCF binding, intracellular NOTCH1 transcriptional complexes recognize RBPJ, the DNA binding sequence motif, and recruit SWI/SNF / BAF complexes. **b**, With gained CTCF binding, NOTCH1, BAF complexes and CTCF protein molecules cooperatively alter the chromosome conformation and form a transcriptional condensate (dashed circle) to regulate expression of the target gene.

Our work in T-ALL found that gains in CTCF binding are located in distal enhancer regions, while cancer-specific CTCF binding loss events are enriched at gene promoter regions and correlate with repressed transcription of these promoters and decreased chromatin interactions. Recently, an enhancer-docking mechanism described by Schuijers et al. [62] proposed that a single CTCF binding upstream of a promoter can function as a docking site for multiple distal enhancers; in this way, multiple enhancers loop to a single CTCF site to activate a single target gene promoter[62]. Loss of such a docking CTCF site then removes the ability to form these multiple enhancer loops, thus greatly reducing the ability to activate transcription. While our observations of cancer-specific lost CTCF sites are consistent with this “enhancer docking” model, further studies are required to understand the causal relationships between CTCF binding loss and gene repression.

Our study is built upon integrative computational analyses of multi-source public data coupled with our multi-omics experimental validations using T-ALL as a model system. As a pan-cancer study, our work is limited by data availability and quality. For example, coverage of RRBS and sequencing depths might lead to potential underestimation of differential DNA methylation. Nevertheless, existence of gained CTCF binding independent of DNA methylation is validated. Our findings pave the way for further mechanistic studies of causal relationships between CTCF binding alteration and oncogenic TF activities in leukemia as well as other cancers. Following our proposed model, oncogenic drivers can lead to novel CTCF binding at distinct enhancer regions in the genome, thus creating a signature pattern of CTCF binding. Having observed evidence supporting this model in T-ALL, we believe that studying aberrant CTCF binding events in other cancer types can further our understanding of the underlying oncogenic transcriptional regulatory networks specific to that cancer. In conclusion, unique aberrant CTCF binding pattern represents a novel epigenomic signature of cancer that can be independent of mutations and DNA methylation changes. Our work provides insights into a new angle of mechanistic research on cancer epigenomics.

## Methods

### EXPERIMENTAL PROCEDURE

#### Patient Xenografting and Cell Culture

Jurkat and CUTLL1 cells have been described previously[36, 63]. Cells were cultured in RPMI1640 medium with L-glutamine and 25mM HEPES (Corning) supplemented with 10% heat-inactivated fetal bovine serum (Sigma-Aldrich), 10 U/mL of penicillin-streptomycin (Gibco), and 1x glutaMAX (Gibco) in a humidified incubator at 37°C and 5% CO_2_. Human CD4^+^ T cells were purchased from AllCells. Primary human samples were collected by collaborating institutions with informed consent and analyzed under the supervision of the Institutional Review Board of Padova University, the Associazone Italiana Ematologia Oncologia Pediatrica, and the Berlin-Frankfurt-Münster (AIEOP-BFM) ALL 2000/2006 pediatric clinical trials. Informed consent to use leftover material for research purposes was obtained from all of the patients at trial entry in accordance with the Declaration of Helsinki.

#### Antibodies and Reagents

Western blots were performed using the following antibodies: Actin and CTCF from Millipore Sigma (clone C4; 07-729) and cleaved NOTCH1 (Val1744) from Cell Signaling Technology (4147). ChIP-seq were performed using the following antibodies: CTCF from Millipore Sigma (07-729), H3K27Ac (8173S) and H3K27me3 (9733S) from Cell Signaling Technology, and H3K4me1 (07-473) from Millipore.

#### In Situ Hi-C

In situ Hi-C was performed on CD4+ T cells, Jurkat, CUTLL1, and patient xenografts as previously described[5]. In brief, cells were crosslinked with 1% formaldehyde for 10 minutes at room temperature. Per Hi-C reaction, 5 million cells were lysed and nuclei were permeabilized. DNA was digested with MboI from New England Biolabs (R0147M). Digested fragments were labeled with biotinylated d-ATP from Jena Bioscience (NU-835-BIO14-S) and ligated. After RNase treatment and Proteinase K treatment to reverse crosslinks, nuclei were sonicated using a Covaris E220 to produce an average fragment length of 400 bp. Streptavidin beads from ThermoFisher Scientific (65001) were used to pull down biotin-labeled fragments. Following purification and isolation of DNA, final libraries were prepared using the NEBNext® Ultra™ II DNA Library Prep Kit for Illumina® and sequenced via paired end sequencing at a read length of 150 bp on an Illumina HiSeq 2500 to produce on average 400 million reads per sample.

#### ChIP-seq Profiling

CD4+ T cells, Jurkat, CUTLL1, and patient xenografts were crosslinked with 1% formaldehyde and 1% fetal bovine serum in PBS for 10 minutes at room temperature. The reaction was quenched with 0.2M glycine at room temperature for 5 minutes. Cells were then washed with PBS and pelleted.

For CTCF ChIPs, immunoprecipitation was performed based on a protocol described previously[64]. A pellet containing 50 million cells was lysed with 5mL of lysis buffer (50 mM HEPES-KOH, pH 7.5, 140 mM NaCl, 1 mM EDTA, 10% glycerol, 0.5% NP-40, 0.25% Triton X-100) for 10 minutes at 4°C. Nuclei were pelleted at 1350xg for 7 minutes and resuspended in 10 mM Tris pH 8, 1 mM EDTA and 0.1% SDS. Chromatin was sheared with a Covaris E220 system to an average fragment length of 400 bp and spun at 15,000 rpm for 10 minutes to remove insoluble chromatin and debris. The supernatant was incubated with 20 μL of Dynabeads Protein G for 30 minutes before discarding the beads. 1% of the total volume was saved as input and the rest was incubated with anti-CTCF antibody overnight. 100 μL of Dynabeads Protein G was added for 2 hours. Bound fragments were washed twice with 1 mL of low salt buffer (20 mM Tris-HCl pH 8.0, 150 mM NaCl, 2 mM EDTA, 1% w/v Triton X-100, and 0.1% w/v SDS), once with high salt buffer (20 mM Tris-HCl pH 8.0, 500 mM NaCl, 2 mM EDTA, 1% w/v Triton X-100, and 0.1% w/v SDS), once with lithium chloride buffer (10 mM Tris-HCl pH 8.0, 250 mM LiCl, 1 mM EDTA, 1% w/v NP-40, and 1% w/v deoxycholic acid) and twice with TE (10mM Tris pH 8, 1 mM EDTA).

For histone ChIPs, cells were lysed in 375 μL of nuclei incubation buffer (15 mM Tris pH 7.5, 60 mM KCl, 150 mM NaCl, 15 mM MgCl_2_, 1 mM CaCl_2_, 250 mM sucrose, 0.3% NP-40, 1 mM NaV, 1 mM NaF, and 1 EDTA-free protease inhibitor tablet (Roche)/10 mL in H_2_O) for 10 min on ice. Nuclei were washed once with digest buffer (10 mM NaCl, 10 mM Tris pH 7.5, 3 mM MgCl_2_, 1 mM CaCl_2_, 1 mM NaV, 1 mM NaF, and 1 EDTA-free protease inhibitor tablet (Roche)/10 mL in H_2_O) and resuspended in 57-μL Digest Buffer containing 4.5 units MNase (USB) for 1 hour at 37°C. MNase activity was quenched for 10 minutes on ice upon the addition of EDTA to a final concentration of 20 mM. Nuclei were pelleted and resuspended in 300-μL Nuclei Lysis Buffer (50 mM Tris-HCl pH 8.0, 10 mM EDTA pH 8.0, 1% SDS, 1 mM NaV, 1 mM NaF, and 1 EDTA-free protease inhibitor tablet (Roche)/10 mL in H_2_O) before sonication with a Bioruptor Pico (Diagenode) for 5 minutes (30 seconds on, 30 seconds off). Lysate was centrifuged at max speed for 5 minutes to remove debris. 9 volumes of IP Dilution Buffer (0.01% SDS, 1.1% Triton X-100, 1.2 mM EDTA pH 8.0, 16.7 mM Tris-HCl pH 8.0, 167 mM NaCl, 1 mM NaV, 1 mM NaF, and 1 EDTA-free protease inhibitor tablet (Roche)/10 mL in H_2_O) were added to the supernatant. 50 μL of Dynabeads Protein G were added and the sample was incubated at 4°C for 30 minutes, rotating. 1% of the sample was kept as input, and the remaining sample was split into 3 tubes. 50 μL of Dynabeads Protein G conjugated to 15 μL of the appropriate antibody were added to each tube prior to overnight incubation at 4°C, rotating. Bead-bound complexes were washed for 5 minutes each in 1 mL of low salt buffer, high salt buffer, LiCl buffer, and twice with TE.

To elute bead-bound complexes, beads were resuspended in 50 μL of elution buffer (100 mM NaHCO_3_, 1% w/v SDS) and incubated at 65°C for 15 minutes, shaking at 1,000 RPM on a thermomixer (ThermoScientific). Elution was repeated a second time, and then 100 μL RNase Buffer (12 μL of 5M NaCl, 0.2 μL 30 mg/mL RNase, and 88 μL TE) was added to each ChIP and input sample. Samples were incubated at 37°C for 20 minutes, followed by the addition of 100 μL of proteinase K buffer (2.5 μL 20 mg/mL proteinase K, 5 μL 20% SDS, and 92.5 μL TE) overnight at 65°C. An equal volume of phenol:chloroform solution was added and mixed thoroughly. The mixture was transferred to MaXtract High Density tubes (Qiagen) and centrifuged for 8 minutes at 15,000 rpm. The upper phase was transferred to new tubes and mixed with 1.5 μL 20 mg/mL glycogen, 30 μL 3 M sodium acetate and 800 μL ethanol. Samples were incubated at -80°C until frozen and then centrifuged at 15,000 rpm for 30 minutes at 4°C. The supernatant was removed and pellets were washed in 800 μL 70% ice cold ethanol and spun for 10 minutes at 4°C at 15,000 rpm. Following careful removal of ethanol, pellets were air-dried and resuspended in 30 μL of 10mM Tris at pH 8.

IP and input DNA were then quantified using a Qubit 3.0 fluorometer. Libraries were prepared using the KAPA HyperPrep Kit (KK8505) and sequenced with an Illumina NextSeq 500 to an average depth of 28 million reads per sample.

#### RNA-seq Profiling

RNA was isolated from 3 million cells per sample using the Bio-Rad Aurum™ Total RNA Mini Kit and quantified with the Agilent RNA 6000 Nano Kit with the Agilent Bioanalyzer. Libraries were prepared by rRNA depletion using the Illumina TruSeq® Stranded mRNA Library Prep Kit for a low concentration of starting sample and sequenced by single end sequencing on an Illumina NextSeq 500 to an average depth of 18 million reads per sample.

#### DNA Methylation Profiling

Genomic DNA was isolated using the AllPrep DNA/RNA Micro Kit (Qiagen). To assess genome-wide DNA methylation status, we performed mRRBS (PMCID PMC1258174). Following fluorometric quantification using a Qubit 3.0 instrument, we digested genomic DNA with the restriction enzyme MspI (New England Biolabs) and size selected for fragments approximately 100–250 base pairs in length using solid phase reversible immobilization (SPRI) beads (MagBio Genomics). Resulting DNA underwent bisulfite conversion using the EZ DNA Methylation-Lightning Kit (Zymo Research). We created libraries from bisulfite-converted single stranded DNA using the Pico Methyl-Seq Library Prep Kit (Zymo Research), which were then pooled for sequencing on an Illumina NextSeq 500 instrument using the NextSeq 500/550 V2 High Output reagent kit (1 × 75 cycles) to a minimum read depth of 50 million reads per sample.

#### Whole Genome Sequencing

3 million cells from cell lines or patient samples were pelleted and resuspended in 1 mL of Cell Lysis Solution (Qiagen) mixed with 500 μg of RNase A. The lysis reaction was carried out at 37°C for 15 minutes. 333 μL of Protein Precipitation Solution (Qiagen) was added to each sample which was then vortexed and then centrifuged at 2000 x g for 10 minutes. The supernatant was mixed with 1 mL of isopropanol until DNA strands precipitated from solution. Upon discarding the supernatant, the DNA pellet was washed with 1 mL of 70% ethanol and centrifuged at 2000 x g for 1 minute. The ethanol was then poured out and the pellet was air-dried for 15 minutes before resuspension in 50 to 100 μL of DNA Hydration Solution (Qiagen). DNA was sequenced with paired end Illumina sequencing at 30x coverage.

#### Immunoprecipitation

100 million cells for each immunoprecipitation reaction were pelleted and incubated in Buffer A (10 mM HEPES pH 8.0, 1.5 mM MgCl_2_, 10 mM KCl, 0.5mM DTT) for 10 minutes on ice. Cells were then lysed upon 12 strokes with a 7mL loose pestle tissue grinder (Wheaton, 357542) and centrifuged at 2000 rpm for 7 minutes. Nuclear pellets were resuspended in 5 volumes of TENT buffer (50 mM Tris pH 7.5, 5 mM EDTA, 150 mM NaCl, 1% Triton X-100, 5 mM MgCl_2_) and treated with benzonase for 30 minutes before 5 passages through a 25g x 5/8 in syringe. The insoluble fraction was removed following centrifugation at 2000 rpm for 7 minutes and incubated overnight with Dynabeads Protein G hybridized with antibody. 2 million cells were removed for input. Beads and nuclei lysates were washed 6 times with TENT buffer and then eluted in 0.1M glycine pH 2.5 with 100 mM Tris pH 8.0 prior. NuPAGE LDS sample buffer was added to eluates and inputs, which were then incubated at 70°C for 15 minutes before analysis by western blot.

### PUBLIC DATA COLLECTION

Public CTCF ChIP-seq data were collected from Cistrome Data Browser[65] (for peak files) and NCBI GEO[66] (for fastq files, Table S1). Histone modification ChIP-seq data were collected from NCBI GEO and ENCODE[67] (for bam files). Public RNA-seq data in multiple cell types were collected from ENCODE (for fastq files). DNA methylation profiling data were collected from ENCODE (for bed bedMethyl files) and NCBI GEO. Hi-C data were collected from NCBI GEO and ENCODE (for fastq files). ATAC-seq data were collected from NCBI GEO (for fastq files). Whole Genome Sequencing data for BRCA, COAD, LUAD and PRAD samples were collected from International Cancer Genome Consortium (ICGC) Data Portal[47]. Survival data for BRCA, COAD and LUAD were downloaded from the TCGA Data Portal[68]. Detailed information including accession IDs of all public datasets collected in this work can be found in Table S4.

### DATA PROCESSING

#### ChIP-seq Data Analysis

Sequence alignment for ChIP-seq data in fastq files was performed using the same standard analysis pipeline as used in Cistrome DB[65], for consistence and reproducibility. All sequence data genomic alignment were performed using the Chilin[69] pipeline with default parameters ($ chilin simple -p narrow [--pe] -s hg38 --threads 8 -t IN.fq -i PRENAME -o OUTDIR). Briefly, sequence reads were aligned to the human reference genome (GRCH38/hg38) using BWA[70] ($ bwa aln -q 5 -l 32 -k 2 -t 8 INDEX IN.fq > PRENAME.sai $ bwa {samse | sampe} INDEX PRENAME.sai IN.fq > PRENAME.sam). Sam files were then converted into bam files using samtools[71] ($ samtools view -bS -q 1 -@ 8 PRENAME.sam > PRENAME.bam). For CTCF ChIP-seq datasets, MACS2[72] was used to call peaks under the FDR threshold of 0.01 ($ macs2 callpeak --SPMR -B -q 0.01 --keep-dup 1 -g hs -t PRENAME.bam -n PRENAME -- outidr OUTDIR). Peaks with fold enrichment of at least 4 were retained. Bigwiggle files were generated using BEDTools[73] and UCSC tools[74] ($ bedtools slop -i PRENAME.bdg -g CHROMSIZE -b 0|bedClip stdin CHROMSIZE PRENAME.bdg.clip $ LC_COLLATE=C sort -k1,1 -k2,2n PRENAME.bdg.clip > PRENAME.bdg.sort.clip $ bedGraphToBigWig PRENAME.bdg.sort.clip CHROMSIZE PRENAME.bw). Finally, only the CTCF ChIP-seq samples that have at least 2,000 peaks were included in the downstream integrative analysis.

#### ATAC-seq Data Analysis

Trim Galore[75] was used to trim the raw sequencing reads ($ trim_galore --nextera --phred33 -- fastqc --paired R1.fq R2.fq -o OUTDIR). Reads were aligned to the human reference genome (GRCH38/hg38) using Bowtie2[76] ($ bowtie2 -p 10 -X 2000 -x INDEX -1 R1.fq -2 R2.fq -S PRENAME.sam). Sam files were then converted into bam files using samtools[71] ($ samtools view -bS -q 1 -@ 8 PRENAME.sam > PRENAME.bam). Bedtools was used to convert bam files into bed format ($ bamToBed -i PRENAME.bam -bedpe > PRENAME_PE.bed). Reads mapped to mitochondria DNA were discarded from downstream analysis.

#### RNA-seq Data Analysis

RNA-seq datasets were processed using Salmon[77] ($ salmon quant --gcBias -i INDEX -l A -p 8 {-1 R1.fq -2 R2.fq| -r IN.fq} -o OUTDIR). Transcriptome index was built on the human reference genome (GRCH38/hg38). Transcripts-level abundance estimates were summarized to the gene-level using the “tximport”[78] package for differential expression analysis. DESeq2[79] was used to identify differentially expressed genes, and different thresholds used in different analysis were listed correspondingly in the manuscript.

#### Hi-C Data Analysis

Hi-C data were processed using HiC-Pro[80] ($ HiC-Pro -i INDIR -o OUTDIR -c CONFIG -p). Contact maps were generated at a resolution of 5kb. Raw matrix data were normalized using the approach described in *Normalization of Chromatin Interactions*.

#### DNA Methylation Data Analysis

DNA methylation data (for T-ALL cell lines and T-ALL patients) were demultiplexed with bcl2fastq followed by trimming of 10 base pairs from the 5’ end to remove primer and adaptor sequences using TrimGalore[75]. Sequence alignment to the GRCh38/hg38 reference genome and methylation calls were performed with Bismark[81] ($ bismark --multicore 8 --bowtie2 -q -N 1 INDEX INFILE.fq). Coverage (counts) files for cytosines in CpG context were generated using Bismark[81, 82] ($ bismark_methylation_extractor --multicore 8 --comprehensive --bedGraph INFILE_bismark_bt2.bam).

#### Whole Genome Sequencing Data Analysis

Mutations were identified for two T-ALL cell lines (Jurkat and CUTLL1) and two T-ALL patient samples from the whole genome sequencing data. We aligned the Illumina short-read sequences to the human reference genome (GRCH38/hg38) using BWA[70] mem. We used SAMBlaster[83] to identify the discordant pairs, split reads and flag the putative PCR duplicates. We used SAMBAMBA[84] to convert the SAM aligned into the BAM format, and samtools[71] was used to sort those aligned to create a BAM file corresponding to each sample.

We used VarDict[85] to identify the variants that overlapped the union CTCF binding sites. We used all the default parameters except ‘-f 0.1’ which was used to identify variants that were supported by greater than 10% of the reads at that location. We annotated the variants using Variant Effect Predictor (VEP)[86], and used custom scripts to identify the variants that influence TF binding.

We again used VarDict[85] to identify the variants in the CTCF and NOTCH1 genes for the four samples. We used all the default parameters except ‘-f 0.1’ which was used to identify variants that were supported by greater than 10% of the reads at that location. We annotated the variants using Variant Effect Predictor (VEP)[86], and then filtered it to identify the mutations that were either (a) not seen in more than 1% of any normal human population, or (b) had a CADD score of deleteriousness > 20, or (c) was present in the COSMIC database.

### INTEGRATIVE MODELING AND STATISTICAL ANALYSIS

#### Identification of CTCF Binding Repertoire in the Human Genome

For CTCF ChIP-seq, we included a total of 793 datasets, including 787 public datasets and 6 datasets we generated (Table S1). 765 public CTCF ChIP-seq datasets with peaks more than 2,000 were further included in this study. Each dataset can yield MACS2-identified CTCF peaks in the range between 2,050 and 198,021, with a median of 46,451, and a total of 36,873,077 peaks (Fig. S1b). The distribution of the interval lengths between adjacent CTCF peak summits of all 36,873,077 peaks from 771 datasets has an inflection point at ∼150bp (Fig. S1c) indicating the boundary between the same binding site and different binding sites[87]. Therefore, we used 150 bps as the cutoff to merge CTCF peaks. In practice, we extended +/-75 bps from each peak summit to generate a 150bp region centered at the summit to represent each peak, and merged all overlapping peak regions to generate a union set of CTCF binding sites, which contains 688,429 non-overlapping sites. Each binding site was assigned a CTCF occupancy score, defined as the tally of ChIP-seq datasets that exhibit a peak within the site. Accordingly, we defined the occupancy frequency as the ratio of the occupancy score over the total number of CTCF ChIP-seq datasets. To further ensuring the robustness of the identified CTCF binding sites, we selected 285,467 high-confidence sites with occupancy score ≥3 for downstream analyses. CTCF motifs within the union binding sites were searched by FIMO[88] with Jaspar[89] matrix (ID: MA0139.1), with a p-value threshold of 1e-4. One motif with the smallest p-value was retained for each CTCF binding site.

#### Identification of Constitutive CTCF Binding Sites

The distribution of occupancy scores of all 285,467 CTCF binding sites (Fig. S1d, blue curve) shows that the majority of the CTCF binding sites occur in only a few datasets, and the number of binding sites decreases with increasing occupancy score when the occupancy score is small. However, there are CTCF binding sites that are highly conserved across almost all datasets (e.g., binding sites with occupancy score greater than 600). We use a power law function to fit the distribution curve (blue) shown in Fig. S1d to determine the cutoff for constitutive CTCF sites. We denote 0*_i_* as the number of observed CTCF binding sites with occupancy score equal to *i*, and *E_i_* as the number of expected CTCF sites with occupancy score equal to *i*. The power law fitting to data 0*_i_* can be described as (Fig. S1d, green):

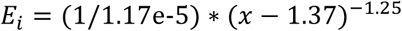

where *x* is the occupancy score. We define the cutoff *A* for constitutive CTCF binding sites as:

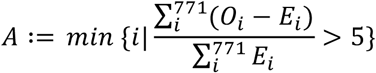

In other words, the total observed CTCF sites with occupancy score greater than *A* should be 6 times more than expected. We then determined *A*=615, and used an occupancy frequency cutoff of 80% to define 22,097 constitutive CTCF binding sites, which corresponds to the occupancy score ≥616 in all 771 CTCF ChIP-seq datasets.

#### Identification of Cancer Specific Gained/Lost CTCF Binding Sites

We used the following 2 criteria to identify cancer-specific lost CTCF binding sites: 1) The CTCF binding site should have a significantly lower occupancy frequency for datasets of that cancer type compared to the occupancy frequency for all datasets; and 2) CTCF binding level (quantified as normalized ChIP-seq read counts) at the site is significantly lower in cancer datasets than in other datasets. For gained CTCF sites, we used the *vice versa* set of criteria. Briefly, for each CTCF binding site in each cancer type, the occupancy score in the cancer datasets were calculated along with its occupancy score in all 771 datasets. CTCF binding levels were obtained from a normalized read count matrix in which the ChIP-seq read counts (RPKM) were first calculated for union CTCF binding sites in all datasets and then followed by quantile normalization. We used unpaired two-tailed Student’s *t*-test to quantify the difference of binding levels between different groups of datasets, and the p-value was then adjusted using the Benjamini-Hochberg Procedure[90]. In addition, binding occupancy scores and binding levels were compared between cancer datasets and datasets from the matched normal tissue or cell types, in order to take into account the potential confounding factor of tissue-specificity rather than cancer-specificity. Detailed criteria for identifying cancer specific CTCF binding sites are described below:

- *Cancer-specific lost CTCF binding sites*: (1) occupancy frequency ≤0.2 in cancer datasets; (2) occupancy frequency ≥0.7 in 771 datasets; (3) occupancy frequency ≥0.5 (with occupancy score ≥2) in matched normal tissue datasets; (4) CTCF levels are lower in cancer compared to all other datasets (statistic score <0), (5) CTCF levels are lower in cancer compared to matched normal tissue datasets (statistic score <0), (6) averaged CTCF binding signals (RPKM) <5 in cancer datasets.
- *Cancer-specific gained CTCF binding sites*: (1) occupancy frequency ≥0.5 (with occupancy score ≥2) in cancer datasets; (2) occupancy frequency ≤0.2 in 771 datasets; 1. (3) occupancy score =0 in matched normal tissue datasets; (4) CTCF levels are significantly higher in cancer compared to all other datasets (FDR ≤0.01), (5) CTCF binding levels are significantly higher in cancer compared to matched normal tissue datasets (FDR ≤0.01), (6) averaged CTCF binding signals (RPKM) >2 in cancer datasets.

The specific gained and lost CTCF binding sites for each cancer type are shown in Table S3.

#### Quantification of Differential Chromatin Accessibility

We used the processed data from Ref. [40] that include a matrix of normalized ATAC-seq insertion counts within the TCGA pan-cancer peak set to assess the differential chromatin accessibility around CTCF binding sites. For each cancer type among BRCA, CRC, LUAD and PRAD, the pan-cancer ATAC-seq peaks that overlap with identified cancer-type-specific lost or gained CTCF binding sites were used for downstream analyses. The ATAC-seq differential score for each peak was quantified as the fold change of the average of the normalized ATAC-seq insertion counts from patient samples in the corresponding cancer type versus from patients in other cancer types, and the ATAC-seq differential score was then assigned to the peak overlapped CTCF binding site.

For consistency, we applied the same approach used for TCGA ATAC-seq data to analyze the collected ATAC-seq data from T-ALL cell line Jurkat and normal CD4+ T cells. A data matrix was generated using ATAC-seq raw read counts on union CTCF binding sites for all Jurkat and T cells datasets. Quantile normalization was applied on the log2 scaled matrix (pseudo count =5). The ATAC-seq differential score was measured as the fold change of the averaged normalized ATAC-seq counts between datasets of Jurkat versus CD4+ T cell at each CTCF binding site.

#### Survival Analysis

Survival analysis was applied on patient samples having both supported TCGA clinical data and ATAC-seq data[40]. For each cancer type among BRCA, CRC and LUAD, the patient samples are separated into two equal-sized groups based on the chromatin accessibility at identified cancer-specific lost or gained CTCF binding sites as follows: (1) For each cancer type, get the pan-cancer ATAC-seq peaks that also present in cancer-specific lost or gained CTCF binding sites. (2) Rank the patient samples by the normalized ATAC-seq insertion counts at each individual ATAC-seq peak, and assign the sum of ranks across all ATAC-seq peaks to each TCGA patient sample. (3) Separate the patient samples from the corresponding cancer type into two equal-sized groups (top 50% and bottom 50%) based on the sum of all ranks, e.g., patient samples with more or less chromatin accessibility on cancer specific CTCF binding sites. The Kaplan-Meier (K-M) method was then used to create the survival plots and log-rank test was used to compare the differences of survival curves.

#### Normalization of Chromatin Interactions

Given a Hi-C contact map *A* = {a*_ij_*}, the score *a_ij_* reflects mapped reads between two genomic regions *i* and *j.* Suppose the bin size is 5kb, regions *i* and *j* will have a genomic distance of *|i* -*j*|* *5kb*. Since the contact probability between two bins decreases with increasing genomic distance[91], we normalized the contact map as follows: for any given genomic distance *d_k_*= *k* * *5kb,* we quantify a normalization factor 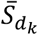 as the averaged interactions among all bin pairs with the same genomic distance *dk* in a same chromosome, e.g., 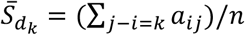, where *n* is the total number of bin pairs with distance *dk.* The interaction score *a_ij_* between two bins with distance *dk* was then normalized by 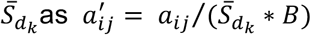, where Bis the number of valid interaction pairs per billion for each sample. Using this approach, we normalized the matrix *A* into 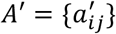 within each chromosome.

#### Detection of Differential Chromatin Interactions

We denoted the normalized Hi-C contact maps in the cancer dataset and the normal dataset as *C* = {*c_ij_*} and *N* = {*n_ij_*}, respectively. For a given CTCF binding site *x* (with coordinate *x_c_)* and a pre-defined genomic distance *L,* the chromatin interactions between *x* and its nearby non-overlapped 5kb bins with genomic distance up to *Lare* collected from *C* and *N* respectively. Specifically, interaction scores between *x* and its nearby 5kb bins in Care collected as *IC* ={*c_ij_*}, while either *i* or *j* equals to 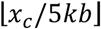, and 0 < *(j* - *i)* * *5kb≤ L.* Similarly, the interaction scores between *x* and its nearby 5kb bins in *N* were collected as *IN* = *{n_ij_}.* A paired two-tailed

Student’s t-test was then applied on *IC* and *IN* to quantify the differential interaction between cancer and normal cells surrounding CTCF binding site *x*.

#### Association of CTCF Binding with Gene Expression

54 cell types for which both CTCF ChIP-seq data and RNA-seq data are publicly available were selected (Table S4) for investigating the association between CTCF binding and gene expression for each CTCF-gene pair in the same chromosome. To obtain the CTCF binding level, a read count matrix was generated using reads per kilobase per million (RPKM) on union CTCF binding sites from ChIP-seq data. The read count matrix was scaled with square root of RPKM followed by quantile normalization. Gene expression level was measured for each gene using the square root of transcripts per million (TPM) from RNA-seq data. For each CTCF-gene pair, we quantified the association between the CTCF site and the gene across all 54 cell types using the correlation coefficient *R* between the normalized CTCF binding level and gene expression (Fig. 2e). CTCF-gene pairs were deemed “highly correlated” with *R*^2^ greater than 0.25.

#### Identification of Constitutive CTCF-Bounded Chromatin Domains

For each CTCF binding site, we defined its associated chromatin domain as the genomic region that: (1) includes this specific CTCF binding site; (2) is bounded by a pair of constitutive CTCF binding sites with motifs of opposite orientations; and (3) occupy a minimum of 100kb and a maximum of 1MB region on each side of the CTCF binding site. Fig. 2f contains schematic of how constitutive CTCF-bounded chromatin domains were defined.

#### Detection of DNA Methylation Changes Surrounding CTCF Binding Sites

DNA methylation changes were detected within a 300bp region centered at each CTCF binding site. Regions with at least 3 CpGs covered by at least 5 reads (≥5x) in both cancer cell lines and matched normal tissues were retained. A 300bp region was detected as differentially methylated if the averaged differential methylation levels of all CpGs (≥5x) within this region was greater than 20%[92].

#### Detection of Differential Mutation Rate and Motif Score

For each CTCF binding site, the mutation rate was calculated as the occurrence of mutation events against the number of samples/patients at each single base pair within a 400bp region centered at the CTCF binding site. The mutation rate for a group of CTCF binding sites was considered as the averaged mutation rate for each base pair within the 400bp region.

Motif score was measured by scoring the CTCF position weight matrix (Jaspar[89], Matrix ID: MA0139.1) to a 19bp DNA sequence centered at the CTCF motif or CTCF binding site using log likelihood ratios (with background nucleotide frequency as [0.275,0.225, 0.225, 0.275] for A,C,G,T). The differential motif score was calculated by comparing motif scores for the reference and the mutated sequences.

#### Identification of CTCF Intra-Domain Differentially Interacted Regions

For a given set of CTCF binding sites, the chromatin interaction changes between a CTCF site and each of its intra-domain non-overlapped bins, measured from normalized Hi-C contact maps in cancer cells over matched normal cells, were collected for each of the CTCF binding sites (Fig. S11b). Regions with decreased interactions (log2 FC <-1, averaged log2 interaction >0) with cancer-specific lost CTCF binding sites, and regions with increased interactions (log2 FC >1, averaged log2 interaction >0) with cancer-specific gained CTCF binding sites were used for downstream transcription factor (TF) enrichment analysis.

#### Transcription Factor Enrichment Analysis

A revised version of the BART algorithm[51] was used for TF enrichment analysis. Briefly, a collection of union DNase I hypersensitive sites[93] (UDHS) was previously curated as a repertoire of all candidate cis-regulatory elements in the human genome, and 7032 ChIP-seq datasets were collected for 883 TFs[51], with each TF having one or more ChIP-seq datasets from multiple cell types or conditions. A binary profile was generated for each TF on UDHS indicating whether the TF has at least one peak from any of its ChIP-seq datasets locate within each of the UDHS. Binding enrichment analysis was applied for each TF by comparing the TF binding on a subset of UDHS overlapping the selected genomic regions versus the TF binding on UDHS. *P* value was obtained using two-tailed Fisher’s exact test.

## Declarations

### Availability of data and materials

The datasets generated in this study are available in NCBI GEO repository, under accession number GSE130140 (https://www.ncbi.nlm.nih.gov/geo/query/acc.cgi?acc=GSE130140). The public data used and analyzed in this study are summarized in Methods, with the accession information included in Tables S1, 2, and 4. All source code for analyzing the data and generating the results and figures in this paper are available at GitHub (https://github.com/zanglab/CTCF_T-ALL_code).

### Authors’ contributions

C.F., Z.W., P.N., and C.Z. designed the experiments, interpreted the results, and wrote the manuscript. C.Z. and P.N. conceived and directed the study. C.F., C.H., and P.N. performed most experiments. Z.W., A.R., and C.Z. processed the data and conducted computational analyses. B.D.S, K.A.H., E.R.A. and M.E.F. designed, executed and interpreted DNA methylation experiments. S.L.S. and A.G.-M. performed and helped with the design of ATAC-seq experiments. K.P.E. helped with the design, execution and interpretation of Hi-C experiments. All authors read and approved the manuscript.

## Acknowledgements

The authors would like to thank Dr. Jon C. Aster, Dr. Warren S. Pear, and members of the Zang and Ntziachristos laboratories for helpful discussions and critical reading of the manuscript.

