## Supplementary data for "Cancer-specific CTCF binding facilitates oncogenic transcriptional dysregulation"

Supplementary Data includes 11 Supplementary figures and 5 Supplementary tables.

**Figure S1.** Identification of cancer specific CTCF binding sites.

**Figure S2.** Characterization of cancer-specific gained/lost CTCF binding sites: chromatin accessibility and clinical outcome.

**Figure S3.** Cancer-specific lost/gained CTCF binding sites associated with changed local chromatin interactions in different scales.

**Figure S4.** Histone modification patterns at cancer-specific lost/gained CTCF binding sites.

**Figure S5.** Histone modification patterns in normal $CD4^+$ T-cell, T-ALL cell lines and T-ALL patients at $T\text{-}ALL_{lost}$ and $T\text{-}ALL_{gained}$ CTCF binding sites.

**Figure S6.** Cancer-specific loss/gain of CTCF events correlate with gene expression.

**Figure S7.** DNA methylation changes near $T\text{-}ALL_{lost}$ and $T\text{-}ALL_{gained}$ CTCF binding sites in two T-ALL patients.

**Figure S8.** CTCF binding loss/gain events in T-ALL cell lines and T-ALL patients do not associate with DNA sequence mutations.

**Figure S9.** CTCF binding loss/gain events in 5 cancer types do not associate with DNA sequence mutations observed in ICGC samples.

**Figure S10.** Cancer specific gained CTCF correlate with oncogenic transcription factor.

**Figure S11.** Cancer specific gained CTCF binding sites correlate with oncogenic transcriptional activation.

Supplementary tables are available upon request.

**Table S1.** List of collected CTCF ChIP-seq datasets.

**Table S2.** Lists of CTCF ChIP-seq datasets in cancer cell lines and corresponding normal tissues for identification of cancer-specific CTCF binding sites.

**Table S3.** Lists of cancer-specific lost and gained CTCF binding sites in six cancer types.

**Table S4.** Lists of collected public multi-omics data, including ATAC-seq, DNA methylation, Hi-C, ChIP-seq and RNA-seq used in this study.

**Table S5.** BART prediction results for oncogenic TFs in T-ALL and CRC.

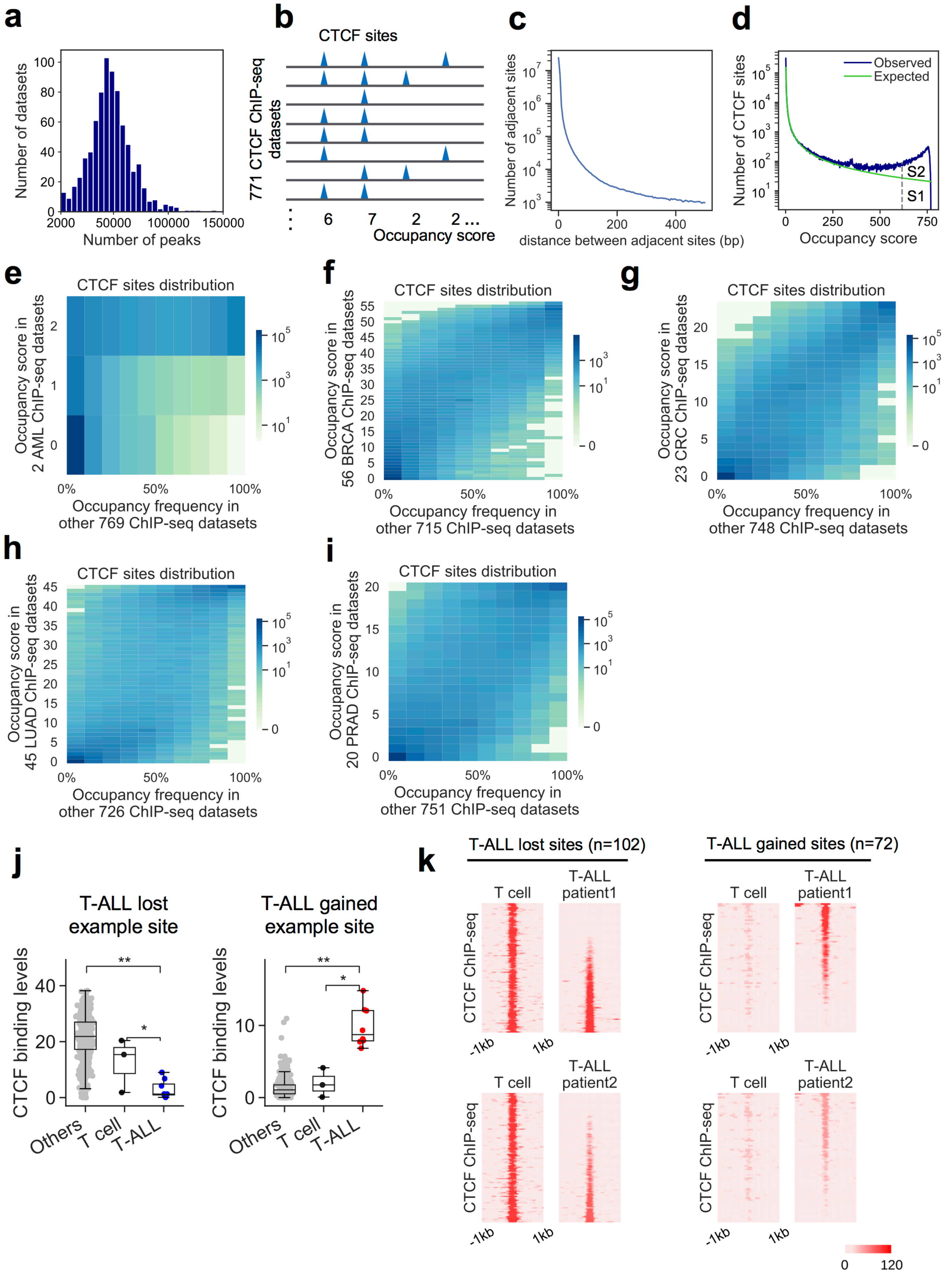
a
Number of datasets
Number of peaks
b
CTCF sites
771 CTCF ChIP-seq datasets
6 7 2 2 ...
Occupancy score
c
Number of adjacent sites
distance between adjacent sites (bp)
d
Number of CTCF sites
Observed
Expected
S2
S1
Occupancy score
e
CTCF sites distribution
Occupancy score in 2 AML ChIP-seq datasets
Occupancy frequency in other 769 ChIP-seq datasets
f
CTCF sites distribution
Occupancy score in 56 BRCA ChIP-seq datasets
Occupancy frequency in other 715 ChIP-seq datasets
g
CTCF sites distribution
Occupancy score in 23 CRC ChIP-seq datasets
Occupancy frequency in other 748 ChIP-seq datasets
h
CTCF sites distribution
Occupancy score in 45 LUAD ChIP-seq datasets
Occupancy frequency in other 726 ChIP-seq datasets
i
CTCF sites distribution
Occupancy score in 20 PRAD ChIP-seq datasets
Occupancy frequency in other 751 ChIP-seq datasets
j
T-ALL lost example site
T-ALL gained example site
CTCF binding levels
Others
T cell
T-ALL
k
T-ALL lost sites (n=102)
T-ALL gained sites (n=72)
T cell
T-ALL patient1
T-ALL patient2
CTCF ChIP-seq
-1kb
1kb
0
120

**Figure S1. Identification of cancer specific CTCF binding sites.**

**a**, Distribution of the number of identified peaks for all 771 CTCF ChIP-seq datasets. Only CTCF ChIP-seq datasets with peaks ≥2000 were included in the downstream study.

**b**, Schematic of integrative analysis of CTCF binding sites from 771 ChIP-seq datasets. An occupancy score was assigned to each union binding site as the tally of ChIP-seq datasets exhibiting a peak within this union binding site region.

**c**, Distribution of the lengths of intervals between adjacent CTCF peak summits. A total of 36,873,077 CTCF peaks were collected from 771 ChIP-seq datasets.

**d**, Distribution of occupancy scores of all 688,429 union CTCF binding sites (blue), and a power law model fitting the distribution (green). The vertical dotted line represents the cutoff of 616 for constitutive CTCF binding sites. S1 represents the number of expected CTCF binding sites with occupancy score more than 616, and S2 represents the number of observed CTCF binding sites with occupancy score more than 616 that exceeding the model-expected.

**e-i**, Distribution of CTCF binding occupancy score in cancer cell lines (y) vs. the CTCF binding occupancy frequency score in the other ChIP-seq datasets (x). Color density in each element represents the number of CTCF binding sites with designated scores. **e**, AML, **f**, breast cancer (BRCA). **g**, colorectal cancer (CRC), **h**, lung cancer (LUAD), **i**, prostate cancer (PRAD).

**j**, Quantile normalized CTCF read counts in normal $CD4^+$ T-cells (black), T-ALL cell lines Jurkat and CUTLL1 (blue for $T\text{-}ALL_{lost}$, red for $T\text{-}ALL_{gained}$) and the other cell types (grey) at a $T\text{-}ALL_{lost}$ site (left) and a $T\text{-}ALL_{gained}$ site (right). *P* values were obtained using unpaired two-tailed Student's *t*-test.

**k**, CTCF ChIP-seq signals at a 2kb region centered at $T\text{-}ALL_{lost}$ (left) and $T\text{-}ALL_{gained}$ (right) CTCF binding site in normal $CD4^+$ T-cells and two T-ALL patient samples.

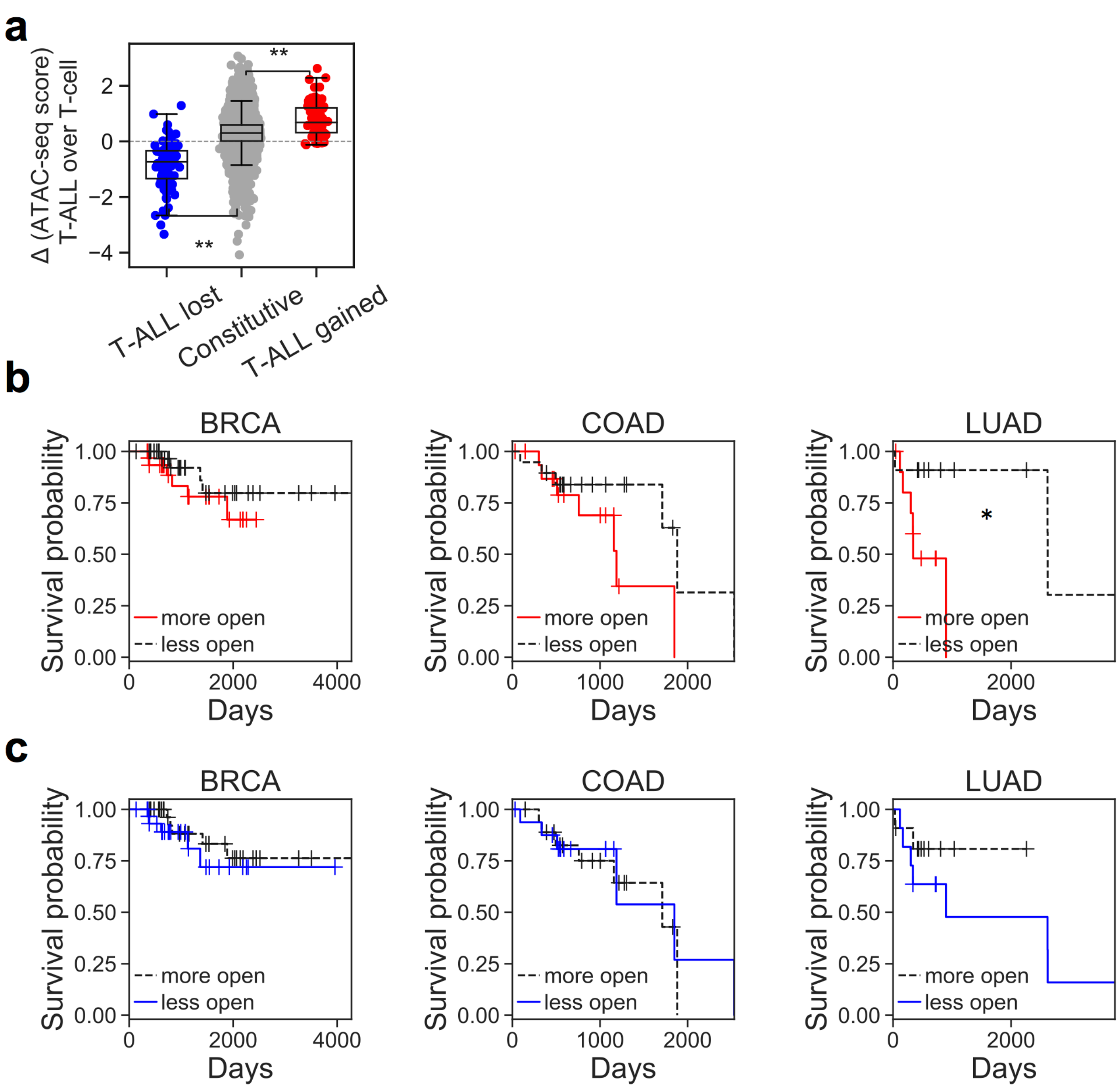


**Figure S2. Characterization of cancer-specific gained/lost CTCF binding sites: chromatin accessibility and clinical outcome.**

**a**, Differential chromatin accessibility (ATAC-seq) in T-ALL cell line Jurkat compared to CD4$^{+}$ normal T-cell at $\textit{T-ALL}_{lost}$ (blue), constitutive (grey) and $\textit{T-ALL}_{gained}$ (red) CTCF binding sites. *, $p<0.05$, **, $p<0.001$, by unpaired two-tailed Student's *t*-test.

**b**, Clinical survival outcomes of TCGA BRCA, COAD and LUAD patients with more open (red) and less open (grey) chromatin accessibility at cancer specific gained CTCF sites. *, $p<0.05$, **, $p<0.001$, by log-rank test.

**c**, Clinical survival outcomes of TCGA BRCA, COAD and LUAD patients with less open (blue) and more open (grey) chromatin accessibility at cancer specific lost CTCF sites.

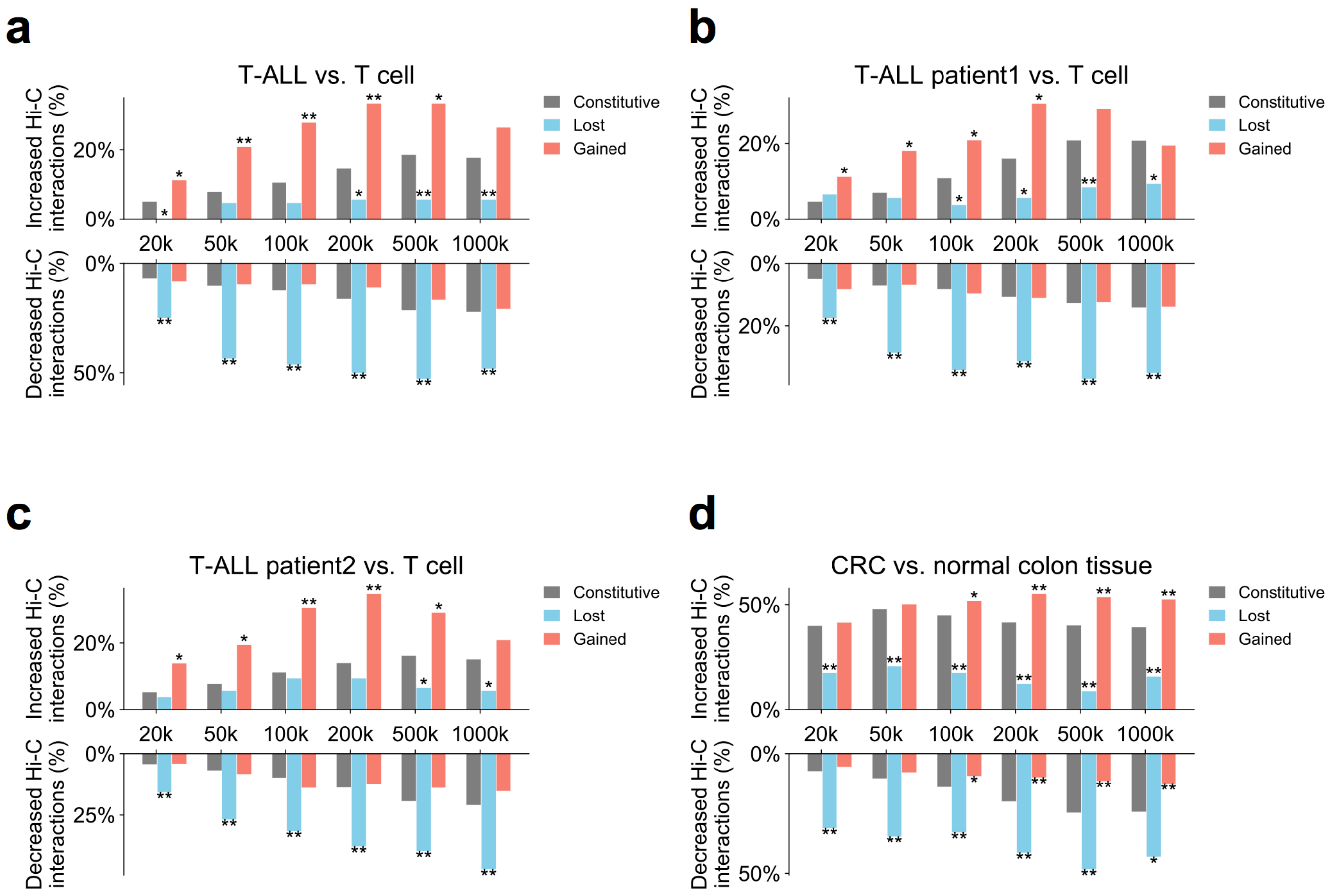


**Figure S3. Cancer-specific lost/gained CTCF binding sites associated with changed local chromatin interactions in different scales.**

**a-d**, Percentage of cancer specific lost (blue), gained (red) and constitutive (grey) CTCF binding sites with increased (top) or decreased (bottom) local chromatin interactions in T-ALL cell line Jurkat (**a**), two T-ALL patients (**b**,**c**) and CRC (**d**) compared to corresponding normal matched tissue as observed in Hi-C. Local chromatin interactions are defined as interactions between a CTCF binding site and 5kb bins located within 20kb, 50kb, 100kb, 200kb, 500kb, 1000kb, respectively, from the CTCF site. *, $p<0.05$, **, $p<0.001$, by two-tailed Fisher's exact test.

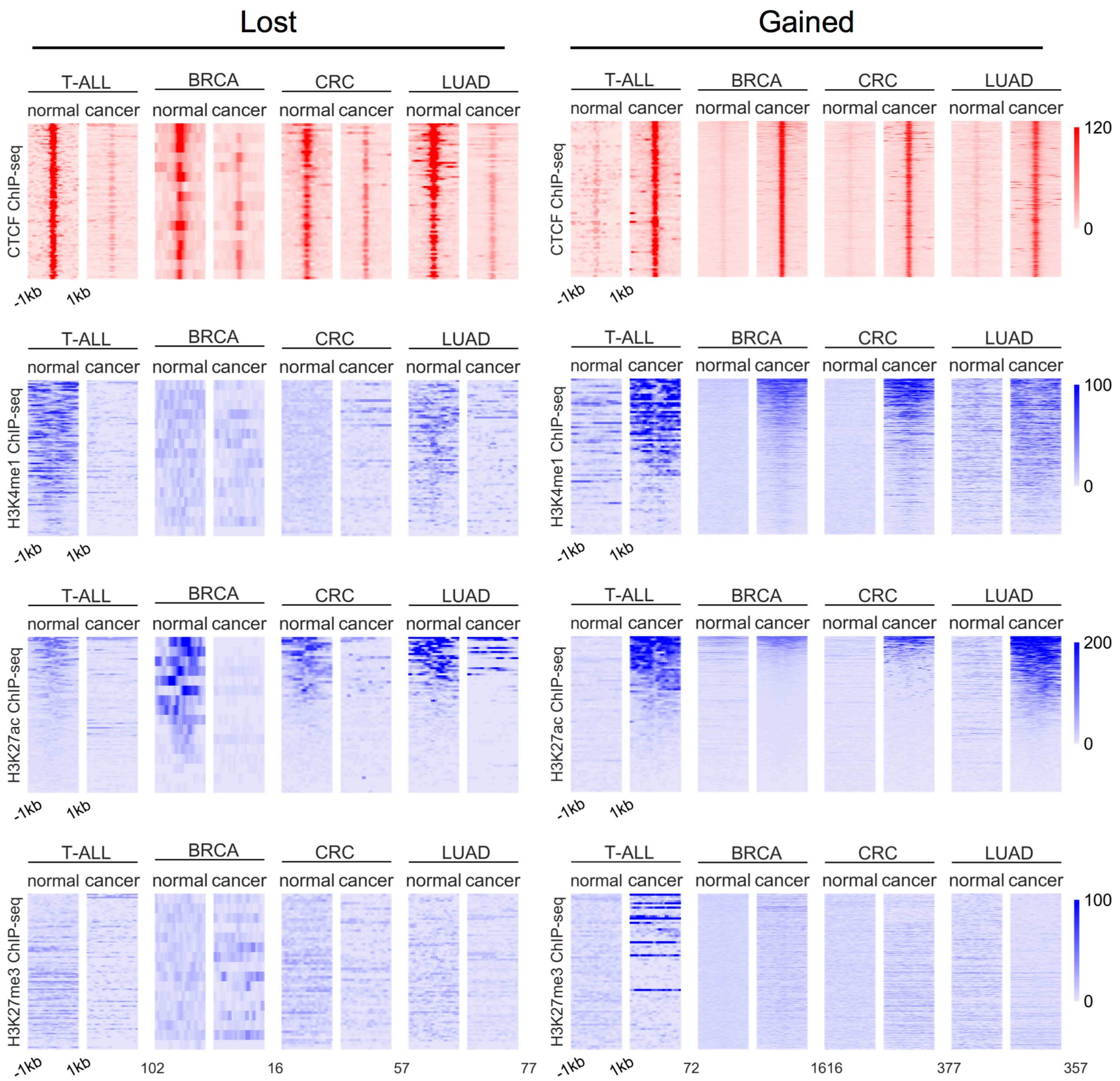


**Figure S4. Histone modification patterns at cancer-specific lost/gained CTCF binding sites.**

Normalized ChIP-seq read counts of CTCF (1st row), H3K4me1 (2nd row), H3K27ac (3rd row), and H3K27me3 (4th row) surrounding identified cancer specific lost (left) and gained (right) CTCF binding sites comparing matched normal tissue and cancer cell lines for T-ALL, BRCA, CRC and LUAD. ChIP-seq heatmaps cover 2kb regions centered at each CTCF site. Rows in corresponding ChIP-seq heatmaps in each cancer type are ranked identically.

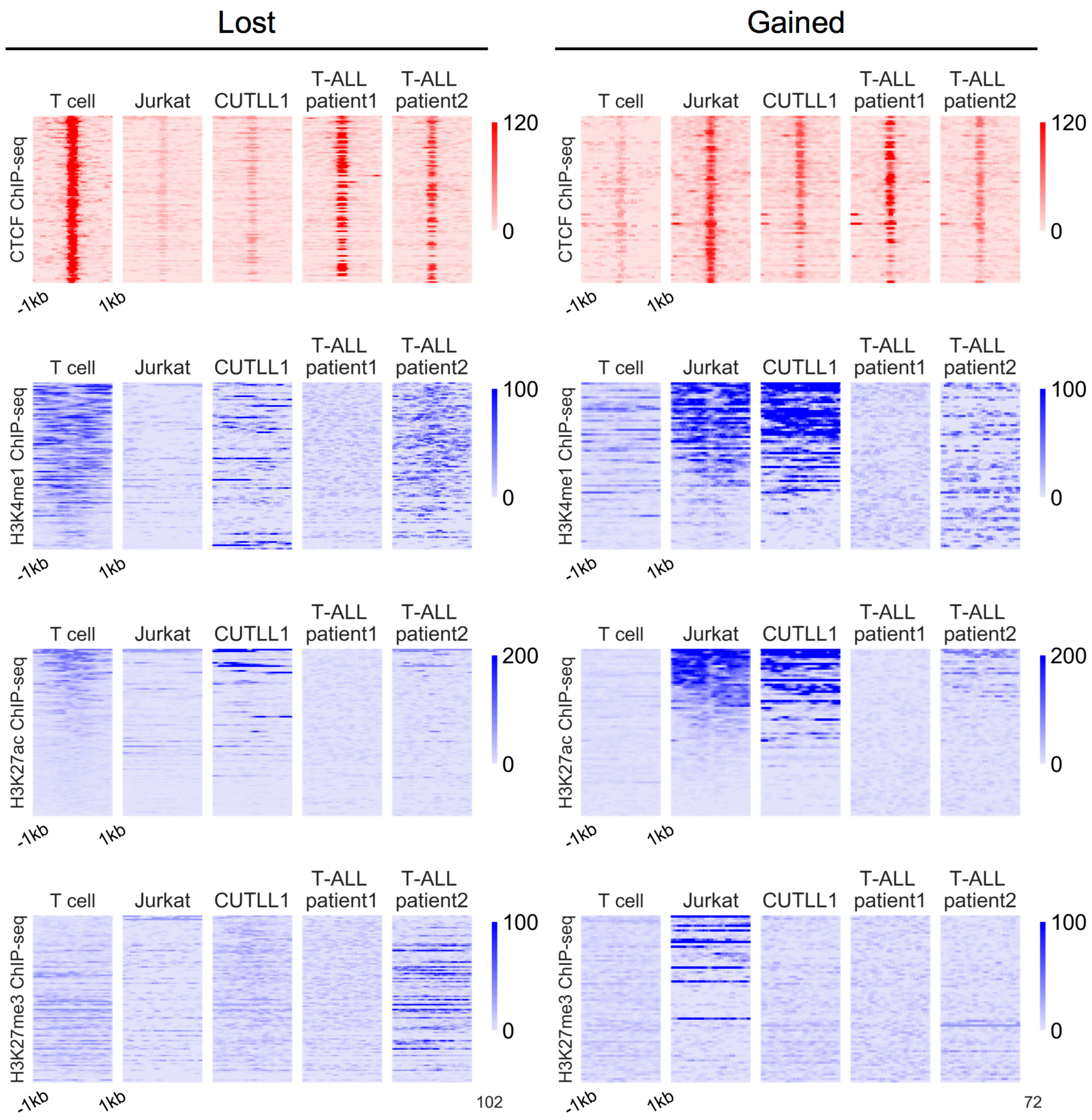


**Figure S5. Histone modification patterns in normal CD4⁺ T-cell, T-ALL cell lines and T-ALL patients at *T-ALL$_{lost}$* and *T-ALL$_{gained}$* CTCF binding sites.**

Normalized ChIP-seq read counts of CTCF (1$^{st}$ row), H3K4me1 (2$^{nd}$ row), H3K27ac (3$^{rd}$ row), and H3K27me3 (4$^{th}$ row) surrounding identified *T-ALL$_{lost}$* (left) and *T-ALL$_{gained}$* (right) CTCF binding sites comparing normal CD4$^{+}$ T-cell, two T-ALL cell lines Jurkat and CUTLL1, and two T-ALL patients. ChIP-seq heatmaps cover 2kb regions centered at each CTCF site. Rows in corresponding ChIP-seq heatmaps are ranked identically.

**a**

| | No. of CTCF | No. of merged region | No. of merged region overlapped w/ HiC boundary | No. of HiC boundary (total 2947) overlapped w/ merged region |
|---|---|---|---|---|
| union | 285,467 | 101 | 53 (52.5%) | 2,946 (100.0%) |
| constitutive | 22,097 | 2,295 | 1,753 (76.4%) | 2,544 (86.3%) |
| boundary | 13,771 | 2,842 | 2,064 (72.6%) | 2,482 (84.2%) |

**b**

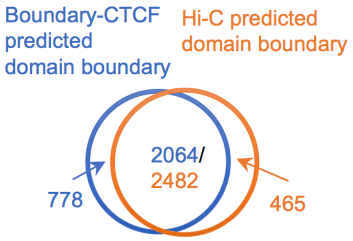


**c**

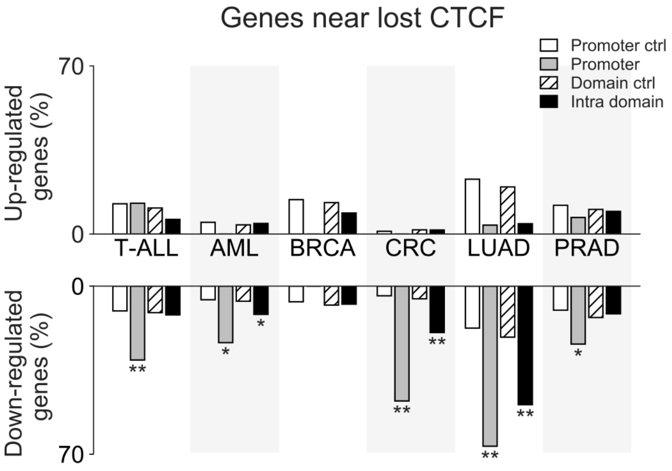


**d**

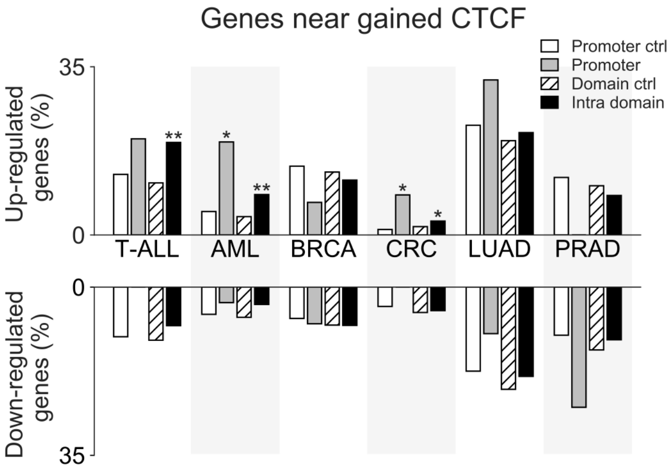


**e**

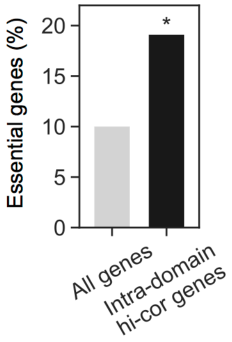

**Figure S6. Cancer-specific loss/gain of CTCF events correlate with gene expression.**

**a,** Comparison of union CTCF, constitutive CTCF and boundary CTCF defined domain boundaries and Hi-C map defined domain boundaries. Boundary CTCF are defined as constitutive CTCFs used as domain boundaries. 40kb extended CTCF sites located within 200kb with each other were merged to generate the merged region in each group.

**b,** Venn diagram comparing boundary CTCF defined domain boundaries and Hi-C map defined domain boundaries.

**c,d**, Percentage of genes that are up-regulated (top, log2FC>1, FDR<1e-5) or down-regulated (bottom, log2FC<-1, FDR<1e-5) located in the chromatin domains containing cancer specific lost (**c**) or gained (**d**) CTCF binding sites in T-ALL, AML, BRCA, CRC, LUAD and PRAD. Promoter: genes having a promoter region (TSS +/-2kb) that contains its paired CTCF binding site; Intra domain: genes paired with a CTCF binding site located with the same chromatin domain. *, p<0.05, **, p<0.001, by two-tailed Fisher's exact test.

**e,** Percentage of genes in each group that are essential genes in T47D cells. Essential genes are defined as genes with lowest β-scores from genome-wide CRISPR screens. Black, genes located in the chromatin domain containing $BRCA_{gained}$ CTCF sites and are highly correlated with the CTCF sites. *, p<0.05, **, p<0.001, by two-tailed Fisher's exact test.

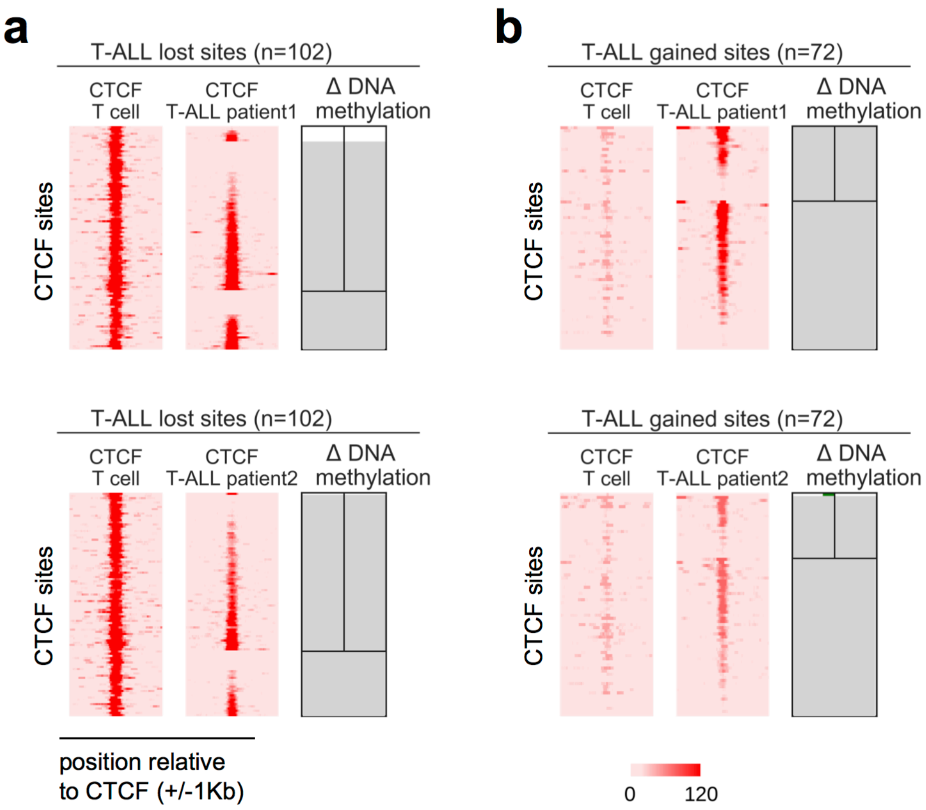


**Figure S7. DNA methylation changes near *T-ALL$_{lost}$* and *T-ALL$_{gained}$* CTCF binding sites in two T-ALL patients.**

**a,b**, ChIP-seq signals and differential DNA methylation levels surrounding *T-ALL$_{lost}$* (**a**) and *T-ALL$_{gained}$* (**b**) CTCF binding sites comparing two T-ALL patients and normal CD4$^+$ T-cell. ChIP-seq heatmaps cover 2kb regions centered at each CTCF site. DNA methylation plots cover 300bp regions centered at each CTCF site. Purple bars represent increased and green bars represent decreased DNA methylation levels (with values in a range from 0 to 100). Grey area with the center vertical line (above the horizontal line) represents regions without enough signal to make confident call of differential methylation. Grey area without the center vertical line (below the horizontal line) represents regions without any detectable methylation signal. Rows in corresponding ChIP-seq and DNA methylation plots are ranked identically.

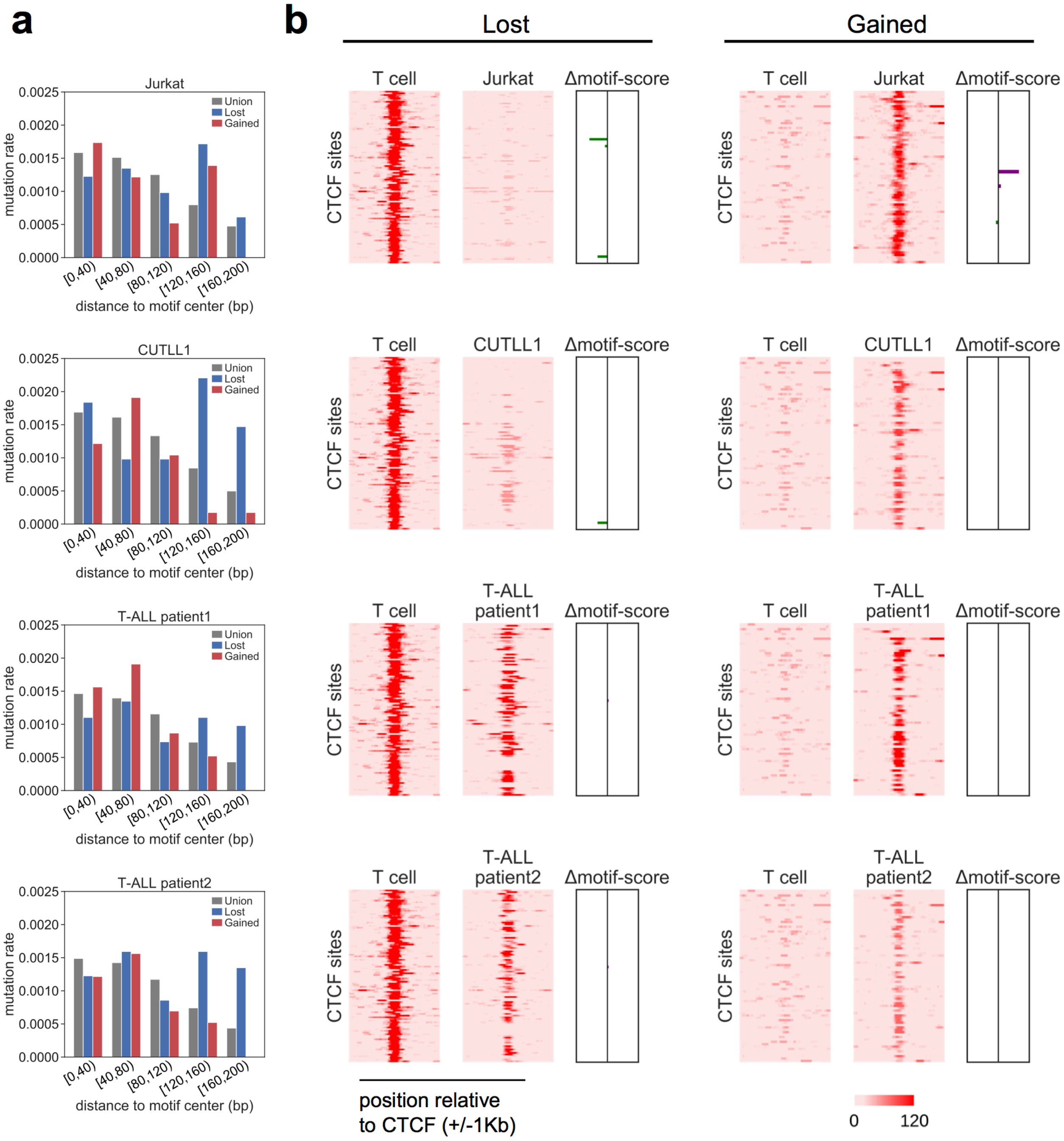
a
b
Lost
Gained
Jurkat
CUTLL1
T-ALL patient1
T-ALL patient2
Union
Lost
Gained
mutation rate
0.0025
0.0020
0.0015
0.0010
0.0005
0.0000
[0,40)
[40,80)
[80,120)
[120,160)
[160,200)
distance to motif center (bp)
T cell
Δmotif-score
CTCF sites
position relative to CTCF (+/-1Kb)
0
120

**Figure S8. CTCF binding loss/gain events in T-ALL cell lines and T-ALL patients do not associate with DNA sequence mutations.**

**a**, Mutation rate in two T-ALL cell lines Jurkat and CUTLL1 and two T-ALL patients surrounding $T\text{-}ALL_{lost}$ (blue), $T\text{-}ALL_{gained}$ (red), and all union (grey) CTCF binding sites. Mutation rate plots cover 400bp regions centered at each CTCF motif and were averaged per 40bp non-overlapped bins at both sides.

**b**, CTCF ChIP-seq signals and motif score changes surrounding $T\text{-}ALL_{lost}$ (left) and $T\text{-}ALL_{gained}$ (right) CTCF sites comparing normal $CD4^{+}$ T-cell, two T-ALL cell lines Jurkat and CUTLL1 and two T-ALL patients. ChIP-seq heatmaps cover 2kb regions centered at each CTCF site. Differential motif score plots cover 19bp CTCF motif sequences. Purple bars represent increased and green bars represent decreased motif score (with values in a range from 0 to 6). Rows in corresponding ChIP-seq heatmaps and differential motif score plots are ranked identically.

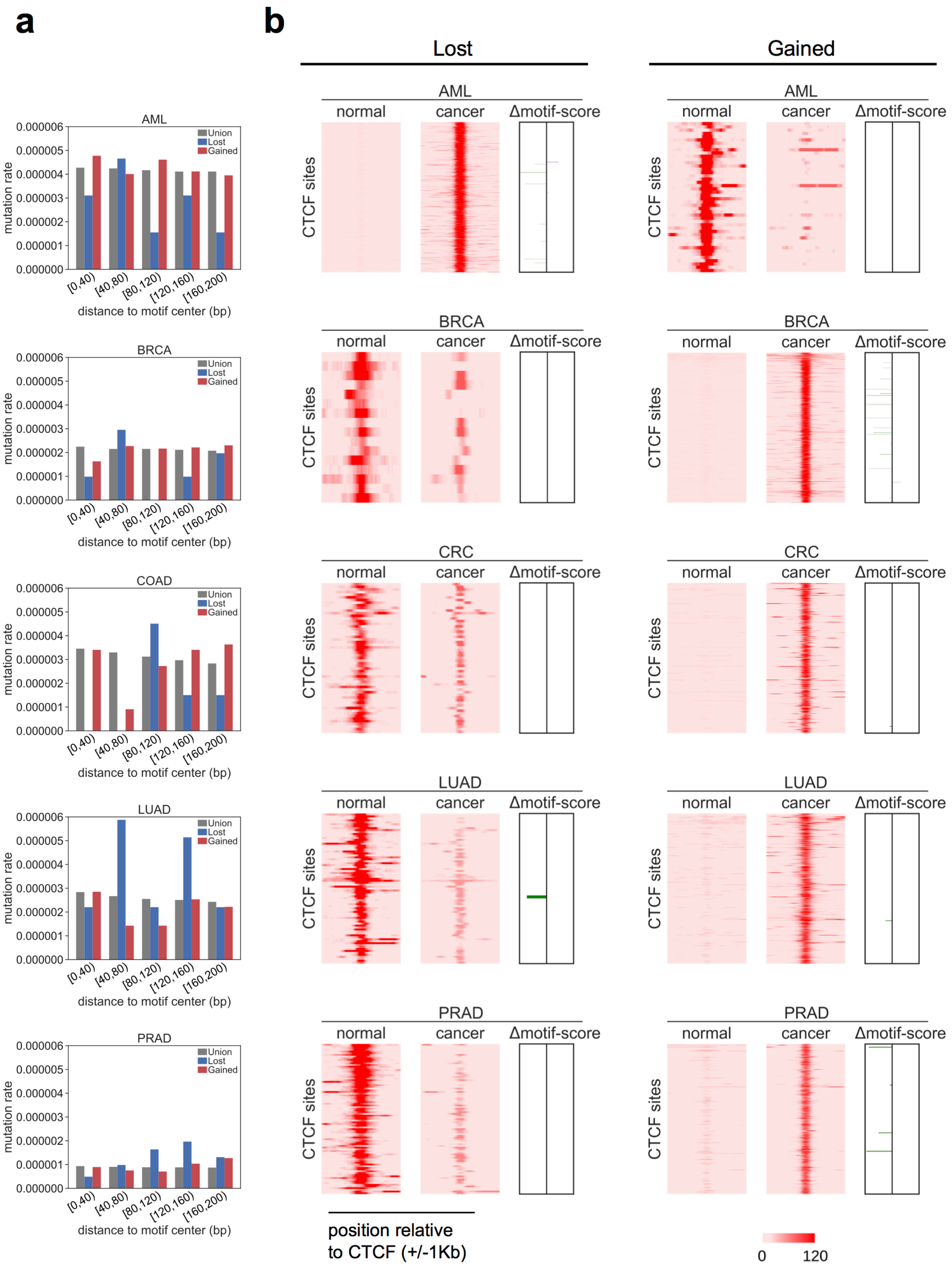

a
b
Lost
Gained
AML
BRCA
COAD
CRC
LUAD
PRAD
Union
Lost
Gained
mutation rate
0.000006
0.000005
0.000004
0.000003
0.000002
0.000001
0.000000
[0,40)
[40,80)
[80,120)
[120,160)
[160,200)
distance to motif center (bp)
normal
cancer
Δmotif-score
CTCF sites
position relative
to CTCF (+/-1Kb)
0
120

**Figure S9. CTCF binding loss/gain events in 5 cancer types do not associate with DNA sequence mutations observed in ICGC samples.**

**a**, Mutation rates in each of the 5 cancers of AML, BRCA, CRC, LUAD and PRAD surrounding cancer specific lost (blue), gained (red), and all union (grey) CTCF binding sites. Mutation rate plots cover 400bp regions centered at each CTCF motif and were averaged per 40bp non-overlapped bins at both sides.

**b**, CTCF ChIP-seq signals and motif score changes surrounding cancer specific lost (left), and gained (right) CTCF sites comparing normal tissues and cancers in each of the 5 cancer types of AML, BRCA, CRC, LUAD and PRAD. ChIP-seq heatmaps cover 2kb regions centered at each CTCF site. Differential motif score plots cover 19bp motif sequences. Purple bars represent increased and green bars represent decreased motif score (with values in a range from 0 to 9). Rows in corresponding ChIP-seq heatmaps and differential motif score plots are ranked identically.

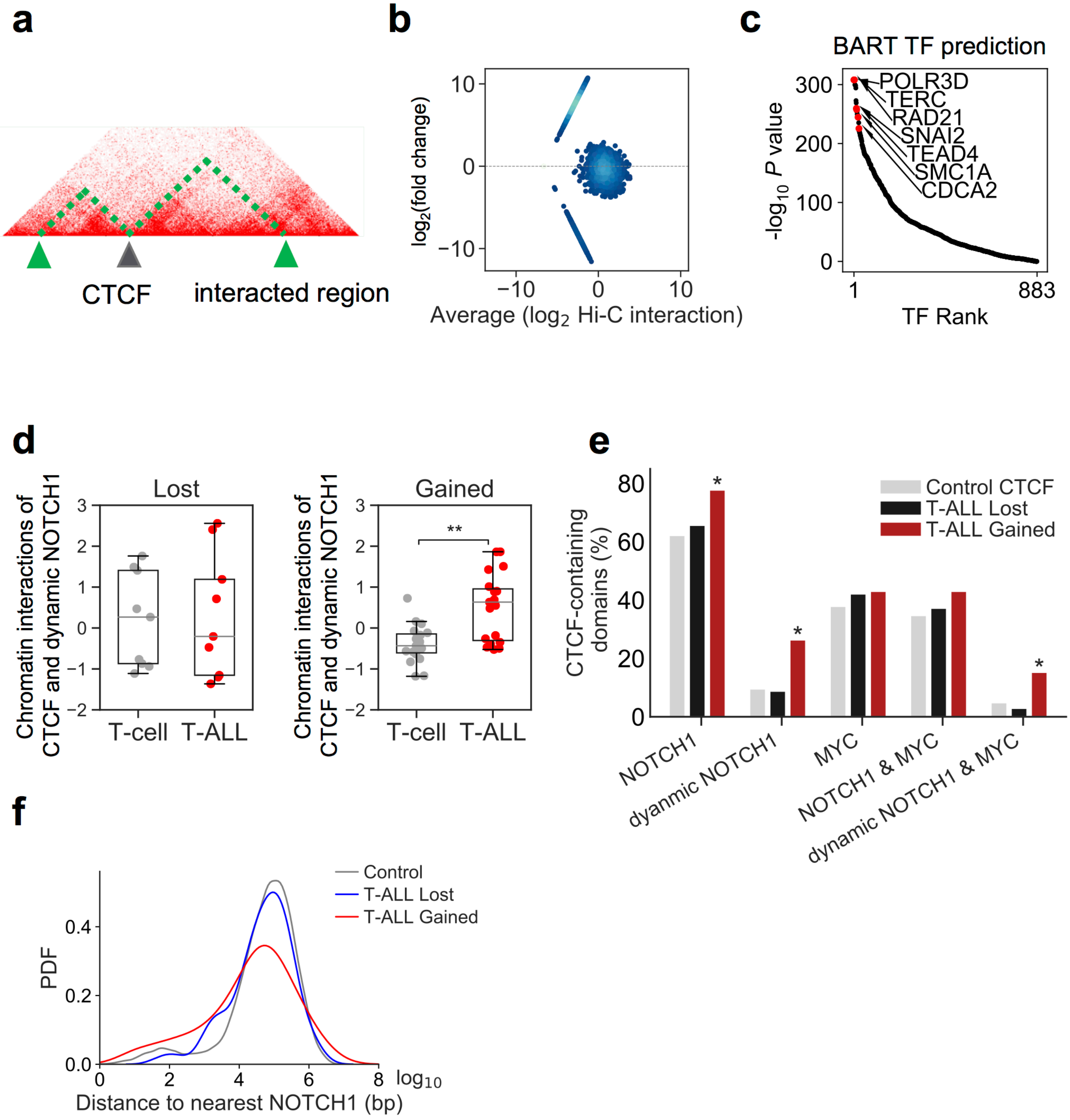
a
CTCF
interacted region
b
log2(fold change)
Average (log2 Hi-C interaction)
c
BART TF prediction
POLR3D
TERC
RAD21
SNAI2
TEAD4
SMC1A
CDCA2
-log10 P value
TF Rank
d
Lost
Gained
Chromatin interactions of CTCF and dynamic NOTCH1
T-cell
T-ALL
**
e
CTCF-containing domains (%)
Control CTCF
T-ALL Lost
T-ALL Gained
NOTCH1
dyanmic NOTCH1
MYC
NOTCH1 & MYC
dynamic NOTCH1 & MYC
f
Control
T-ALL Lost
T-ALL Gained
PDF
log10
Distance to nearest NOTCH1 (bp)

**Figure S10. Cancer specific gained CTCF correlate with oncogenic transcription factor.**

**a,b**, Schematic of identification of cis-domain genomic regions that have changed chromatin interactions with cancer specific gained/lost CTCF sites comparing cancer cell lines and matched normal tissues. **a**, Chromatin interactions between a CTCF binding site and all of its intra-domain 5kb bins. **b**, MA plot showing differential chromatin interactions between cancer and normal cells at cancer specific CTCF binding sites. Each point represents the chromatin interaction changes between a CTCF site and one of its intra-domain 5kb bin.

**c**, BART-predicted transcription factors binding in the genomic regions that have increased interaction with $CRC_{gained}$ CTCF sites comparing HCT116 cell line with normal colon tissue.

**d**, Chromatin interaction levels between $T\text{-}ALL_{lost}$ (left) and $T\text{-}ALL_{gained}$ (right) CTCF binding site and their intra-domain dynamic NOTCH1 binding sites comparing normal $CD4^{+}$ T-cell (grey) and T-ALL cell line CUTLL1 (red) as measured by Hi-C. Chromatin interaction between a CTCF site and a dynamic NOTCH1 site was quantified as a Z-score using interactions between the CTCF site with all of its intra-domain regions as background. *, $p<0.05$, **, $p<0.001$, by paired two-tailed Student's *t*-test.

**e**, Percentage of chromatin domains including different groups of CTCF binding sites that contain a NOTCH1 binding site, a dynamic NOTCH1 binding site, a MYC binding site, a NOTCH1 together with a MYC binding site, and a dynamic NOTCH1 together with a MYC binding site. *, $p<0.05$, **, $p<0.001$, by two-tailed Fisher's exact test.

**f**, Distribution of the distances between the CTCF binding sites in different groups and their nearest NOTCH1 binding sites in CUTLL1 cell line.

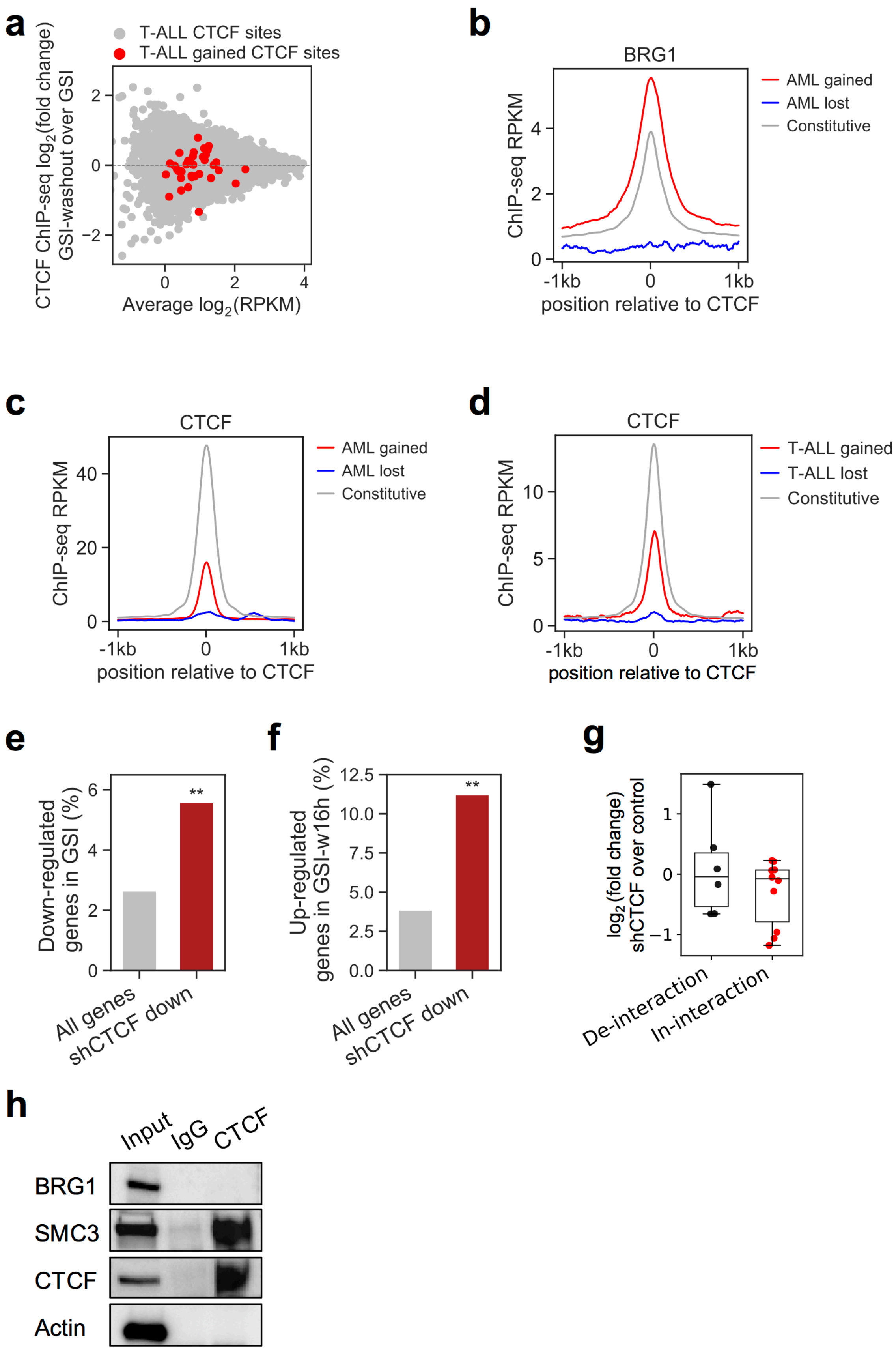

a
T-ALL CTCF sites
T-ALL gained CTCF sites
CTCF ChIP-seq log2(fold change) GSI-washout over GSI
Average log2(RPKM)
b
BRG1
AML gained
AML lost
Constitutive
ChIP-seq RPKM
-1kb
0
1kb
position relative to CTCF
c
CTCF
AML gained
AML lost
Constitutive
ChIP-seq RPKM
position relative to CTCF
d
CTCF
T-ALL gained
T-ALL lost
Constitutive
ChIP-seq RPKM
position relative to CTCF
e
Down-regulated genes in GSI (%)
**
All genes
shCTCF down
f
Up-regulated genes in GSI-w16h (%)
**
All genes
shCTCF down
g
log2(fold change) shCTCF over control
De-interaction
In-interaction
h
Input
IgG
CTCF
BRG1
SMC3
CTCF
Actin

**Figure S11. Cancer specific gained CTCF binding sites correlate with oncogenic transcriptional activation.**

**a**, MA plot showing differential CTCF binding level after GSI wash out treatment in Jurkat. Binding levels of some $T\text{-}ALL_{gained}$ CTCF binding sites (red) are increased.

**b,c** ChIP-seq signals for BRG1 (**b**) and CTCF (**c**) surrounding constitutive (grey), $AML_{lost}$ (blue) and $AML_{gained}$ (red) CTCF binding sites in AML cell line MOLM13. Normalized ChIP-seq read counts (RPKM) covering 2kb regions centered at CTCF binding sites were plotted per 10bp non-overlapped bins.

**d**, ChIP-seq signals for CTCF surrounding constitutive (grey), $T\text{-}ALL_{lost}$ (blue) and $T\text{-}ALL_{gained}$ (red) CTCF binding sites in T-ALL cell line Jurkat. Normalized ChIP-seq read counts (RPKM) covering 2kb regions centered at CTCF binding sites were plotted per 10bp non-overlapped bins.

**e,f**, Down-regulated genes after GSI treatment (**e**) and up-regulated genes after GSI washout treatment (**f**) are enriched for down-regulated genes in shCTCF treatment in CUTLL1. Differentially expressed genes were identified using thresholds of $|log2FC|>0.58$, $FDR<0.01$. *, $p<0.05$, **, $p<0.001$, by two-tailed Fisher's exact test.

**g,** Differential expression upon shCTCF for genes from Group B (fig. 4g, genes located in domains containing both dynamic-NOTCH1 and $T\text{-}ALL_{gained}$ CTCF binding sites) that are up-regulated in CUTLL1 than normal T-cells. Genes were separated if their intra-domain NOTCH1 and $T\text{-}ALL_{gained}$ CTCF sites have decreased (black) or increased (red) chromatin interactions in CUTLL1 compared to normal T-cells.

**h,** CTCF immunopurified proteins from Jurkat cells were resolved on SDS-PAGE gels and interacting partners are visualized by western blot. IgG was immunopurified as a negative control, and SMC3 was immunoblotted as a positive control. IB, immunoblot; IP, immunoprecipitation.
